# Neoproterozoic origin and multiple transitions to macroscopic growth in green seaweeds

**DOI:** 10.1101/668475

**Authors:** Andrea Del Cortona, Christopher J. Jackson, François Bucchini, Michiel Van Bel, Sofie D’hondt, Pavel Škaloud, Charles F. Delwiche, Andrew H. Knoll, John A. Raven, Heroen Verbruggen, Klaas Vandepoele, Olivier De Clerck, Frederik Leliaert

**Affiliations:** Department of Biology, Phycology Research Group, Ghent University, Krijgslaan 281, 9000 Ghent, Belgium; Department of Plant Biotechnology and Bioinformatics, Ghent University, Technologiepark 71, 9052 Zwijnaarde, Belgium; VIB Center for Plant Systems Biology, Technologiepark 71, 9052 Zwijnaarde, Belgium; Bioinformatics Institute Ghent, Ghent University, Technologiepark 71, 9052 Zwijnaarde, Belgium; School of Biosciences, University of Melbourne, Melbourne, Victoria, Australia; Department of Botany, Faculty of Science, Charles University, Benátská 2, CZ-12800 Prague 2, Czech Republic; Department of Cell Biology and Molecular Genetics, University of Maryland, College Park, MD 20742, USA; Department of Organismic and Evolutionary Biology, Harvard University, Cambridge, Massachusetts, 02138, USA; Division of Plant Sciences, University of Dundee at the James Hutton Institute, Dundee, DD2 5DA, UK; School of Biological Sciences, University of Western Australia (M048), 35 Stirling Highway, WA 6009, Australia; Climate Change Cluster, University of Technology, Ultimo, NSW 2006, Australia; Meise Botanic Garden, Nieuwelaan 38, 1860 Meise, Belgium

**Author notes:** K.V, O.D.C and F.L. contributed equally to this work. Author contributions: A.D.C., H.V., K.V., O.D.C, and F.L. designed research; A.D.C., C.J.J., and S.D. performed research; P.S., and C.F.D. contributed cultures and sequence data; A.D.C., C.J.J., F.B., and M.V.B. analyzed data; and A.D.C., A.H.K., J.A.R., H.V., K.V., O.D.C., and F.L. wrote the paper.

**Keywords:** green algae, Chlorophyta, phylogenomics, phylogeny, Ulvophyceae

## Abstract

The Neoproterozoic Era records the transition from a largely bacterial to a predominantly eukaryotic phototrophic world, creating the foundation for the complex benthic ecosystems that have sustained Metazoa from the Ediacaran Period onward. This study focusses on the evolutionary origins of green seaweeds, which play an important ecological role in the benthos of modern sunlit oceans and likely played a crucial part in the evolution of early animals by structuring benthic habitats and providing novel niches. By applying a phylogenomic approach, we resolve deep relationships of the core Chlorophyta (Ulvophyceae or green seaweeds, and freshwater or terrestrial Chlorophyceae and Trebouxiophyceae) and unveil a rapid radiation of Chlorophyceae and the principal lineages of the Ulvophyceae late in the Neoproterozoic Era. Our time-calibrated tree points to an origin and early diversification of green seaweeds in the late Tonian and Cryogenian periods, an interval marked by two global glaciations, with strong consequent changes in the amount of available marine benthic habitat. We hypothesize that the unicellular and simple multicellular ancestors of green seaweeds survived these extreme climate events in isolated refugia, and diversified following recolonization of benthic environments that became increasingly available as sea ice retreated. An increased supply of nutrients and biotic interactions such as grazing pressure has likely triggered the independent evolution of macroscopic growth via different strategies, including both true multicellularity, and multiple types of giant celled forms.

**Significance Statement:** Green seaweeds are important primary producers along coastlines worldwide, and likely played a key role in the evolution of animals. To understand their origin and diversification, we resolve key relationships among extant green algae using a phylotranscriptomic approach. We calibrate our tree using available fossil data, to reconstruct important evolutionary events such as transitions to benthic environments, and evolution of macroscopic growth. We estimate green seaweeds to have originated in the late Tonian/Cryogenian Period, followed by a marked Ordovician diversification of macroscopic forms. This ancient proliferation of green seaweeds likely modified shallow marine ecosystems, which set off an evolutionary arms race between ever larger seaweeds and grazers.

## Introduction

Seaweeds, ecologically important primary producers in marine benthic ecosystems worldwide, have played a prominent role in the global biosphere for many millions of years. Seaweeds comprise red, green, and brown lineages, which evolved independently from unicellular algal ancestors. The red algae are ancient. *Bangiomorpha* and *Raffatazmia*, both reasonably interpreted as red algal fossils, point to an origin of multicellular red algae well into the Mesoproterozoic, 1.0–1.6 Ga ago (1, 2). Brown algae diverged much more recently, with molecular clocks pointing to an Early Jurassic emergence (3, 4). As sister to red algae, microscopic green algae likely originated in Neo- or Mesoproterozoic environments (5, 6), but the lack of a well-supported phylogenetic framework has so far impeded clear interpretation of how many times and when green seaweeds emerged from unicellular ancestors.

There is general consensus that an early split in the evolution of green algae gave rise to two discrete clades. One, the Streptophyta, contains a wide morphological diversity of green algae from freshwater and damp terrestrial habitats, known as charophytes, from which land plants evolved during the Ordovician Period (7, 8). The second clade, the Chlorophyta, diversified as planktonic unicellular organisms, likely in marine as well as freshwater habitats during the late Mesoproterozoic and early Neoproterozoic (9, 10). These ancestral green algae gave rise to several extant lineages of unicellular planktonic marine and freshwater algae, known as the prasinophytes, as well as the core Chlorophyta, which radiated in freshwater, terrestrial, and coastal environments, and evolved a wide diversity of forms ranging from microscopic unicellular and multicellular algae to macroscopic forms (11). Two large core chlorophytan classes, Chlorophyceae and Trebouxiophyceae, are almost entirely restricted to freshwater and terrestrial environments. In contrast, the Ulvophyceae contains the main green seaweed lineages in addition to some smaller microscopic clades, some freshwater species, and the terrestrial Trentepohliales (12, 13). Only two other groups of green seaweeds have phylogenetic affinities outside the Ulvophyceae: the Prasiolales, which belong to the Trebouxiophyceae, and the Palmophyllales, which are allied to prasinophytes (11, 14). These groups are much less diverse compared to the ulvophycean lineages, and clearly evolved independently.

The diversification of Ulvophyceae in marine benthic environments involved the evolution of an astonishing diversity of forms, and most strikingly the evolution of macroscopic, benthic growth forms from small, planktonic unicellular ancestors. Macroscopic growth in green seaweeds presents itself in various forms ranging from multicellular thalli to different types of giant-celled algae with highly specialized cellular and physiological characteristics (15). About ten extant ulvophycean orders are currently recognized, each characterized by a distinctive set of morphological and cytological features (13). Some orders (e.g. Ulvales and Ulotrichales) evolved multicellularity with coupled mitosis and cytokinesis, resulting in uninucleate cells. The Cladophorales evolved siphonocladous multicellular algae, in which mitosis is uncoupled from cytokinesis, resulting in large multinucleate cells with nuclei organized in fixed cytoplasmic domains. The Dasycladales and Bryopsidales evolved siphonous (acellular) macroscopic forms composed of a single giant tubular cell containing thousands to millions of nuclei, or a single macronucleus, and exhibiting cytoplasmic streaming, which enables transport of RNA transcripts, organelles and nutrients throughout the thallus. Some siphonous algae, including species of *Caulerpa*, reach meters in size, thus qualifying as the largest known cells. Although the available data are limited, it seems that acellular algae have similar maximum photosynthetic and nutrient acquisition rates, light absorptance, and ability to growth at low irradiances, to those of morphologically similar multicellular algae (16). The smaller, non-seaweed orders (e.g. Ignatiales, Scotinosphaerales, and Oltmannsiellopsidales), are morphologically less complex, including microscopic unicellular forms with uninucleate cells.

Understanding the origin and ecological diversification of green seaweeds requires a well-resolved phylogeny of the core Chlorophyta, and reliable estimates of the timing of inferred diversification events. Early studies based on ultrastructural features, such as the fine structure the flagellar apparatus, cytokinesis and mitosis have been instrumental in defining higher level groupings of green algae, but they have been inconclusive in determining the relationships among these groups (11). In addition, monophyly of the Ulvophyceae has been questioned because of the absence of shared derived characters (11). Molecular phylogenetic studies based on nuclear and chloroplast gene data often yielded ambivalent or contradictory results (reviewed in 17, 18). These studies make it clear that resolving relationships within the core Chlorophyta is a difficult task, attributed to the antiquity of the clade and possibly further confounded by the rapidity of the early evolutionary radiations (13, 19). An accurate phylogenetic reconstruction will thus require more elaborate sampling, both in terms of species and genes.

Dating divergence times in the phylogeny of the Chlorophyta has been challenging because of difficulties in interpreting fossils with respect to extant taxa. Microfossils in Paleo-to Neoproterozoic rocks have sometimes been assigned to green algae (20-22), but these interpretations are uncertain as they rely on comparisons of simple morphologies (23). Similarly, the assignment of the middle Neoproterozoic filamentous fossil *Proterocladus* (ca. 750 mya) to the Cladophorales (24, 25), is questioned by some (3, 26). Reliable chlorophytan fossils include resistant outer walls of prasinophyte cysts known as phycomata in Ediacaran and Paleozoic deposits (27, 28) and fossils of siphonous seaweeds (Bryopsidales, Dasycladales) from the Cambro-Ordovician onward (29-33). Although reliable green algal fossils from the Neoproterozoic are scarce, organic biomarkers (steroids) indicate that green algae persisted throughout the Cryogenian, and rose to abundance between the Sturtian and Marinoan glaciations (659-645 mya) (34-36).

The principal goal of our study was to resolve the evolutionary relationships among the main lineages of core Chlorophyta and reconstruct key evolutionary events, such as transitions to benthic marine environments, and the evolution of macroscopic growth. We use a rigorous phylotranscriptomic approach, thereby increasing nuclear gene sampling by an order of magnitude and produce a tree calibrated in geological time using available fossil data that facilitates interpreting the ecological and evolutionary context of modern green seaweed origins.

## Results

### Transcriptome data

We collected and analyzed nuclear encoded protein-coding genes from 55 species mined from 15 genomes and 40 transcriptomes (Table S1, S2). 14 transcriptomes were newly generated during this study. Our dataset includes representatives from the major lineages of Streptophyta and Chlorophyta. A denser taxon sampling of the core Chlorophyta, including all main orders of Ulvophyceae (Bryopsidales, Cladophorales, Dasycladales, Ignatiales, Oltmannsiellopsidales, Scotinosphaerales, Trentepohliales, Ulotrichales and Ulvales) was obtained to consolidate evolutionary relationships in this group and advance our understanding of the origin and diversification of green seaweeds.

Eight sequence alignments were assembled for phylogenetic analyses (Table S3). The largest alignment consisted of 539 high-confidence single-copy genes (hereafter referred to as coreGF, see Materials and Methods). From this dataset a subset of 355 genes was selected, with at least one sequence in each ulvophycean order (hereafter referred to as ulvoGF). Partial sequences from the transcriptomes were either scaffolded (scaffolded dataset) or removed (unscaffolded dataset), resulting in a more comprehensive and a more conservative version of the coreGF and ulvoGF datasets (Figure S1). Finally, ambiguously aligned regions were removed to obtain the corresponding trimmed datasets. The eight datasets were analyzed with a range of phylogenetic methods including supermatrix and the coalescence-based analyses. The significance of conflicting topologies was tested to assess the robustness of our findings.

### Green algal phylogeny

Trees estimated by the various methods and datasets were globally well-supported and congruent, be it that different topologies were recovered for a few specific relationships. Most differences were due to analysis method (i.e. supermatrix versus coalescence-based) rather than the dataset used (Figure S3). Chlorodendrophyceae and Pedinophyceae were recovered as the two earliest diverging lineages of the core Chlorophyta, although their relative position differed in the different analyses (Figure 1, Figure S3, Figure S4, Table S4). The Trebouxiophyceae and Chlorophyceae were recovered as monophyletic groups with high support, with the Trebouxiophyceae consisting of two distinct clades, the Chlorellales and the core Trebouxiophyceae. The Trebouxiophyceae were recovered sister to the clade containing the Chlorophyceae and Ulvophyceae in all analyses.

**Figure 1.**
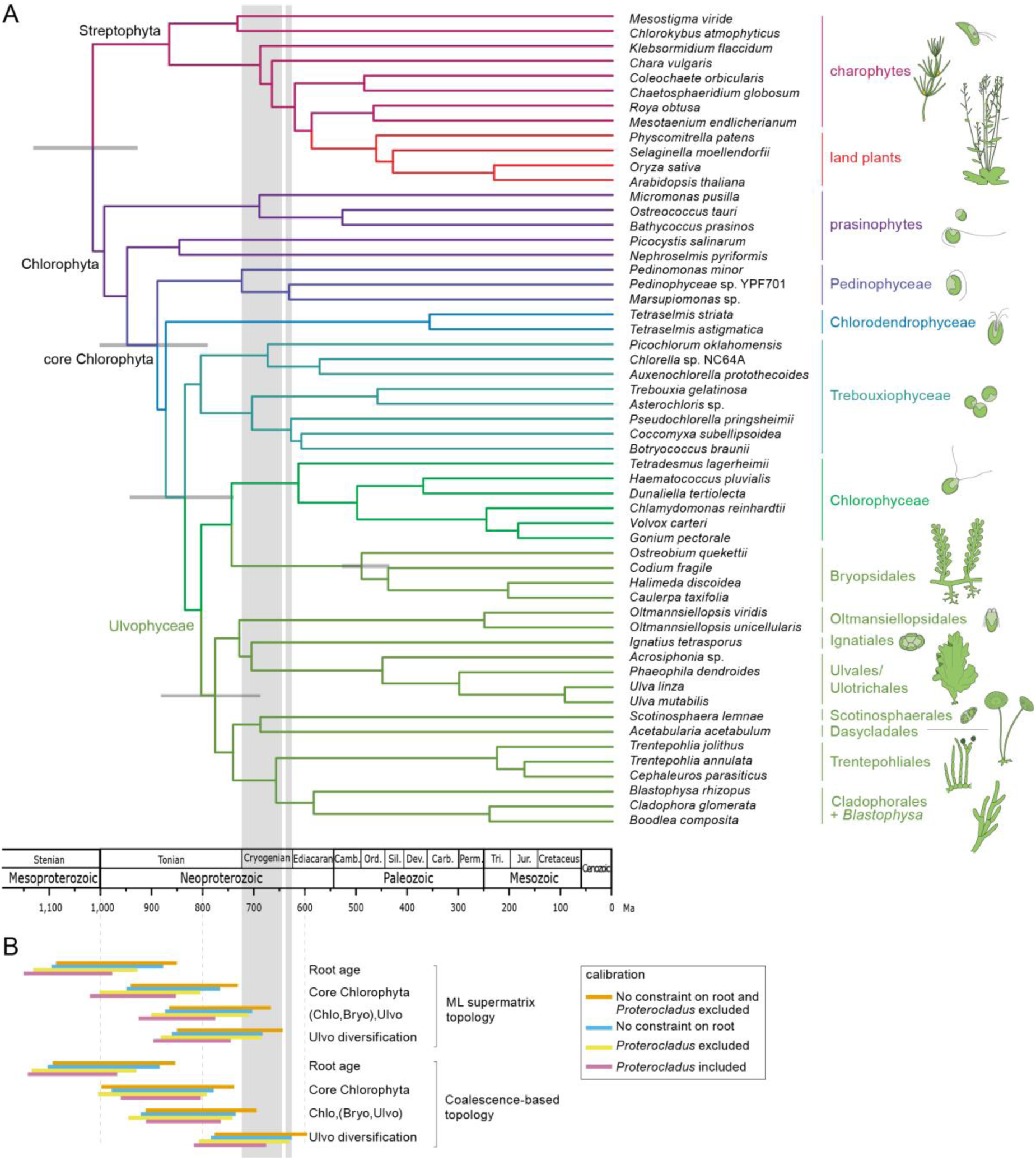
Time-calibrated phylogeny of the green algae. **A**. The topology of the tree is based on the maximum likelihood analysis inferred from a concatenated amino acid alignment of 539 nuclear genes (supermatrix analysis of the coreGF-scaffolded-untrimmed data set, Figure S2). Branch lenghts are based on a relaxed molecular clock analysis of the ten most clock-like genes from the scaffold trimmed dataset, and excluding *Proterocladus* as calibration point. Error bars are indicated for a number of key nodes. **B**. Divergence time confidence intervals of key nodes inferred from different analyses (Figure S7, Table S6).

The siphonous seaweed order Bryopsidales was resolved as the sister clade of the Chlorophyceae, rendering the Ulvophyceae non-monophyletic, in the supermatrix analyses (Figure 1, Figure 2A, Figure S2). Conversely, in the coalescent-based analyses, the Bryopsidales was sister to the remaining Ulvophyceae, with very short branches separating the Bryopsidales, remaining Ulvophyceae, and Chlorophyceae (Figure 2A). A polytomy test could not reject the null hypothesis that the length of the branches in question equals zero, indicating a hard polytomy (Figure 2B, Figure S6). In addition, AU-tests for the trimmed datasets could not reject the topology constrained to conform the coalescence-based analyses (Ulvophyceae monophyletic). For the untrimmed datasets, however, Ulvophyceae monophyly was significantly rejected (Table S4). Analysis of the gene-wise log-likelihood showed that slightly more than half of the genes supported the monophyly of Ulvophyceae for 7 of the 8 datasets (Figure 2C).

**Figure 2.**
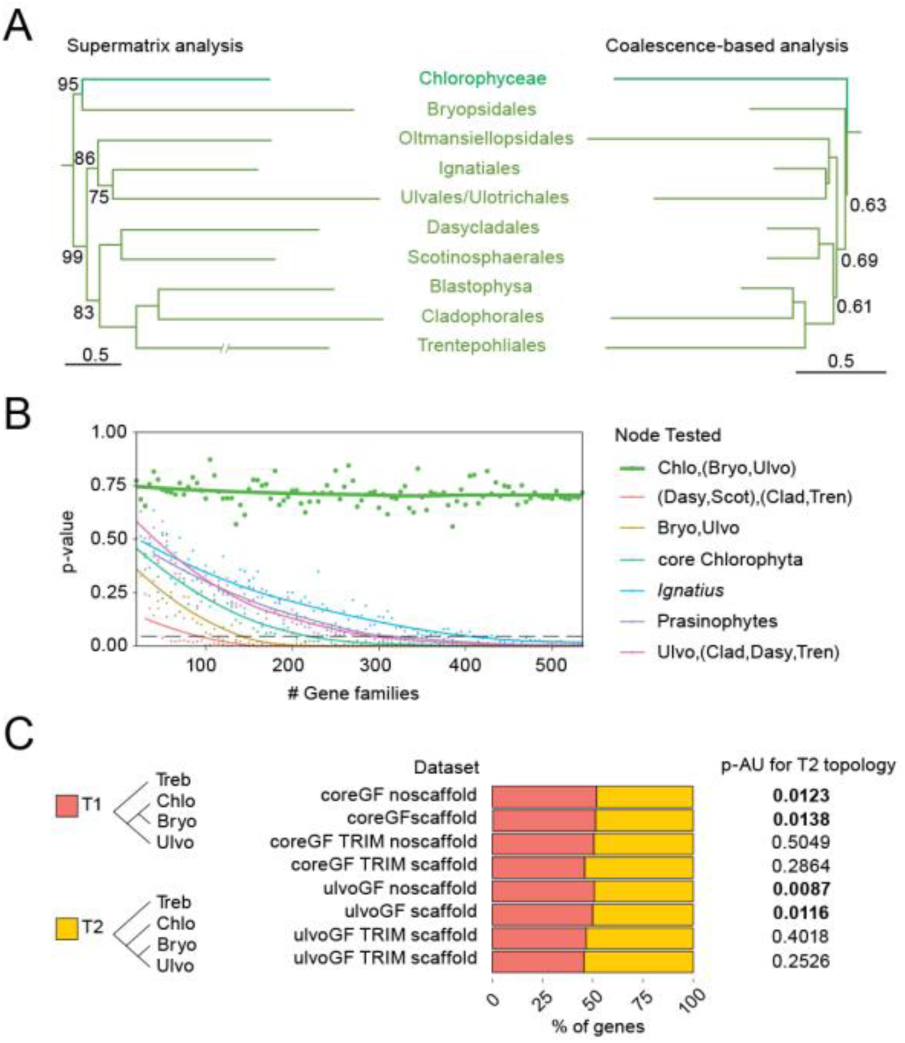
Relationships between Bryopsidales, remaining Ulvophyceae, and Chlorophyceae. **A**. Phylogeny (Chlorophyceae-Ulvophyceae clade pruned) as recovered by the supermatrix (Figure S2) and coalescence-based analysis. **B**. Test for polytomy null-hypothesis for selected relationships based on the coreGF scaffold trimmed dataset (see methods and Figure S6 for details). For the Chlorophyceae, Bryopsidales, and remaining Ulvophyceae (thick green line), increasing gene numbers did not reduce the p-value of rejection. Proportion of genes supporting a sister relationship between Bryopsidales and Chlorophyceae (T1), and a sister relationship between the Bryopsidales and remaining Ulvophyceae (T2), and summary of the AU-test for the constrained topology (T2) (see Table S4 for details). Bryo: Bryopsidales; Chld: Chlorodendrophyceae; Chlo: Chlorophyceae; Clad: Cladophorales+*Blastophysa*; Dasy: Dasycladales; Igna: Ignatiales; Oltm: Oltmannsiellopsidales; Pedi: Pedinophyceae; Scot: Scotinosphaerales; Treb: Trebouxiophyceae; Tren: Trentepohliales; Ulvo: Ulvophyceae (excluding Bryopsidales).

**Figure 3.**
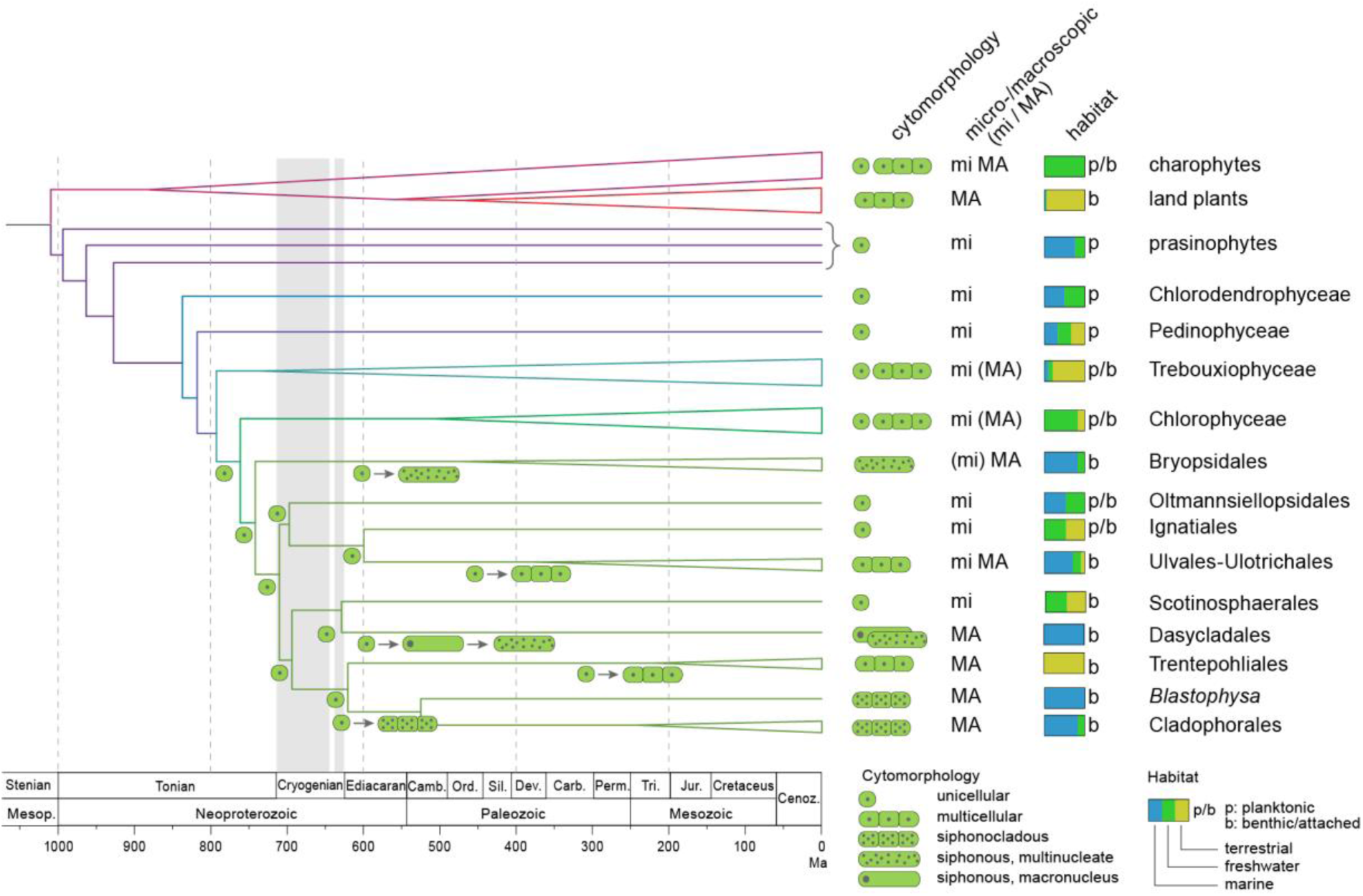
Hypothesis for the evolution of multicellularity and macroscopic growth, and transition to benthic marine habitats in the Ulvophyceae. The topology of the tree is based on the coalescence-based analysis of the coreGF-scaffolded-untrimmed data set. Branch lenghts are based on a relaxed molecular clock analysis of the ten most clock-like genes from the coreGF-scaffolded-trimmed dataset, and excluding *Proterocladus* as calibration point.

The supermatrix and coalescence-based analyses supported the same overall relationships among the remaining orders of Ulvophyceae. Two major clades were recovered, both containing seaweed and unicellular lineages. The position the Ignatiales was not well-supported in any of the phylogenetic analyses, and similar number of genes supported five different relationships (Figure S4, Table S4). The inclusion of *Ignatius* was also found to cause instability in the phylogenetic analyses, and removing it led to an overall higher support for the relationships among the main ulvophycean clades (Figure S5).

### Time-calibrated phylogeny

To estimate the time-frame of diversification, we inferred a chronogram based on 10 of the most clock-like genes, using different molecular clock models, and constraining the analyses with supermatrix- and coalescence-based topologies, respectively. The choice of calibration points largely follows that of Jackson et al. (6). A series of analyses was performed in order to evaluate the effect of calibration nodes (Table S5, Figure S7). Results indicate that the core Chlorophyta emerged during the Neoproterozoic Era, ca. 1000-700 mya (Figure 4B, Table S7). Conditional on the calibration points, diversification of the main ulvophycean lineages took place just before or during the Cryogenian.

### Transition to marine benthic habitats and evolution of macroscopic growth

Our statistical inference of the ancestral states of key ecological and cyto-morphological traits (Figure S8) indicate that the Trebouxiophyceae and Chlorophyceae diversified in freshwater environments mainly. The radiation event at the base of the diversification of Chlorophyceae, Bryopsidales and Ulvophyceae is associated with a major switch back to marine environments early in the evolution of Bryopsidales and Ulvophyceae, populating coastal environments with marine benthic green seaweeds.

## Discussion

We present a phylogenetic reconstruction of the core Chlorophyta based on the most comprehensive multigene dataset to date, and contextualize the relationships in light of transition to marine benthic environments and evolution of multicellularity and macroscopic growth. Although this study focused on the evolution of the Chlorophyta, the relationships among the main lineages of Streptophyta are consistent with current phylogenetic consensus, validating the strength of our phylotranscriptomic approach in confidently resolving difficult phylogenetic relationships, such as identifying the closest living relative of the land plants (37, 38). Our analyses resolve some long-standing phylogenetic questions within the Chlorophyta (39, 40), including the evolutionary placement of the unicellular Pedinophyceae and Chlorodendrophyceae as the earliest diverging lineages of the core Chlorophyta, and monophyly of the Trebouxiophyceae (Supporting information). Our analyses suggest a hard polytomy between the Chlorophyceae, siphonous seaweeds of the Bryopsidales and the remaining Ulvophyceae. A roughly equal numbers of genes supported alternative topologies, indicative of ancient incomplete lineage sorting or a non-bifurcating evolutionary history (41, 42). Although coalescence-based analyses have the ability to detect and incorporate conflicting signals in gene-trees in phylogenetic reconstruction, and supermatrix analyses may converge on a wrong species-tree when incomplete lineage sorting is high (43-45), we hypothesize these lineages represent a rapid ancient radiation.

Our phylogenetic results allow us to propose a scenario for the evolutionary history of green seaweeds. Given that several early-branching ulvophycean lineages consist of unicellular algae (including the Ignatiales, Oltmansiellopsidales, Scotinosphaerales, and some lineages of Ulotrichales and Ulvales), the ancestral ulvophycean was likely a small unicellular organism with a single nucleus. Our phylogeny also affirms the hypothesis by Cocquyt et al. (13) that macroscopic growth evolved independently in various lineages of ulvophyceans from ancestral unicellular green algae, and by different mechanisms: multicellularity with coupled mitosis and cytokinesis (independently in the Ulvales-Ulotrichales and Trentepohliales), multicellularity with uncoupled mitosis and cytokinesis (siphonocladous organization in the Cladophorales and *Blastophysa*) and a siphonous architecture (independently in the Bryopsidales and Dasycladales). The sister relationship of Cladophorales (plus *Blastophysa*) and Trentepohliales may indicate a single origin of multicellularity at the base of this clade, followed by the evolution of multinuclear cells in the Cladophorales. However, the phragmoplast-mediated cell division in Trentepohliales, producing plasmodesmata, differs strongly from cell division in the Cladophorales, which takes place by ingrowth of a diaphragm-like cross wall (11). It may therefore be more reasonable to think that multicellularity evolved independently in the Trentepohliales and the Cladophorales. Contrary to previous phylogenetic studies, which indicated a sister relationship between Bryopsidales and Dasycladales (13, 39), the separate positions of these orders point towards an independent evolution of siphonous organization in Bryopsidales and Dasycladales from ancestral unicellular green algae. The distinct position of the Bryopsidales is unexpected from a morphological view-point, but is supported by ultrastructural features (Supporting information), as well as by independent molecular data. The Trentepohliales, Cladophorales, Dasycladales, and Scotinosphaerales are characterized by a non-canonical nuclear genetic code while the Bryopsidales and all other green algae possess the canonical code (46, 47), pointing towards a single origin of a non-canonical nuclear code in the green algae.

Our relaxed molecular clock analysis indicates that the early diversification of core Chlorophyta took place in the Tonian Period, with several green algal lineages persisting throughout the Cryogenian ice ages. The early diversification of ulvophycean lineages took place during this interval, as well. Diversification before and survival during the Cryogenian glacial interval has been inferred for other groups of eukaryotes based on fossil evidence, including complex forms with blade-stipe-holdfast differentiation that have been interpreted as benthic macroalgae (48-50). It is important to note, however, that the scarce fossil record and the uncertain nature of ancient green algal fossils, along with methodological bias impose a limit on the precision of ancient divergence time estimates (51). These unavoidable uncertainties in node age assignments have to be accommodated in our interpretations, and therefore one cannot dismiss the hypothesis that green seaweeds began to diversify earlier, in the middle or even early Tonian Period. One concrete, well-dated fossil that has been put forward as important to estimate the age of the Ulvophyceae is *Proterocladus*, from ca. 780 Ma shales of the upper Svanbergfjellet Formation in Svalbard, originally interpreted as a *Cladophora*-like alga (25). *Proterocladus* has typical branches that are in cytoplasmic contact with the parent cell, which is as a derived feature in the Cladophorales (52). Assignment of *Proterocladus* to the Cladophorales would imply that the order would be older than 780 my, and would push the divergences of the main ulvophycean lineages even further back in time, inconsistent with our time-calibrated phylogeny, which suggests an Ediacaran or Cambrian origin of the Cladophorales. While *Proterocladus* is certainly eukaryotic, and appears to be coenocytic, we prefer to consider its taxonomic affinity as uncertain based on the late appearance of Cladophorales in our molecular clock analyses.

Ulvophycean diversification could have been promoted by the highly dynamic climatic conditions and habitat variability of the Cryogenian periods (53, 54). Little is known about the paleontological conditions of these periods that would have allowed lineage diversification and survival, but a number of different scenarios can be put forward. The later Tonian Period was a time of evolutionary innovation among both photosynthetic and heterotrophic eukaryotes in a number of major eukaryotic clades (54); thus, inferred divergence of green seaweeds would appear to be part of a broader pattern of Tonian diversification. This has sometimes been related to an increase in atmospheric oxygen levels (e.g., 54), but evidence in support of this hypothesis is mixed, suggesting that any redox change was small relative to later Ediacaran oxygenation (e.g., 55). Alternatively, evolutionary drivers may have included the expansion of eukaryovorous protists in the oceans, supported by both fossils and molecular clock estimates for major eukaryote-eating protistan lineages (56, 57). Experiments show that predation can select for multicellularity in originally unicellular prey populations (58, 59), consistent with the observed and inferred Tonian diversification of multicellular and coenocytic forms, as well as scales and other protistan armor (54, 57).

As already noted, the ensuing Cryogenian Period was characterized by two widespread glaciations, the older, protracted Sturtian ice age and the younger and shorter Marinoan glaciation (53). Among the most extreme climatic events in recorded Earth history, these ice ages, involved the freezing of large parts of the ocean surface for millions of years and dramatic fluctuations in biogeochemical cycling. Snowball Earth conditions were certainly unfavorable for planktonic eukaryotes, resulting in greatly diminished photosynthesis in the marine realm. The fossil record is consistent with this notion, showing low overall eukaryotic diversity during this interval (60). An important key to the success and diversification of ulvophycean green algae during the Cryogenian, thus may have been an evolutionary transition from a planktonic to benthic lifestyle, a shared feature of all extant ulvophycean lineages. During glaciated conditions, benthic environments may have been among the few suitable habitats left for green algal lineages to persist, and diversification may have been promoted in different ways. First, the Sturtian glaciation (716-659 Ma) lasted a long time, but shows evidence of glacial waxing and waning, resulting in secular variation in suitable habitat availability (53). During this period, when colonisable benthic substrates would have oscillated between rare and uncommon, ulvophycean populations likely experienced prolonged isolation and demographic declines that would amplify the effects of genetic drift. We hypothesize that, in concert with local adaptation, this has promoted early diversification of ulvophycean lineages. Macroscopic compressions of probable algae in shales within the Marinoan Nantuo Tillite demonstrate that macroalgae could and did survive at least locally during times when ice was widespread (50). As ice sheets decayed, habitat space would have increased dramatically, perhaps helped along by a transient increase in nutrients (especially P) associated with high post-glacial weathering fluxes. Arguably, the emergence of new ulvophycean lineages could have been rapid in this new permissive ecology, where competition for newly available habitats would have been low, allowing a broader suite of mutations to persist (61).

We hypothesize that following the transition to a greenhouse world at the end of the Marinoan glaciation, surviving lineages would have been able to diversify, and new morphological experiments ensued. The rapid diversification of eukaryotic life in the early Ediacaran Period is evidenced by molecular biomarkers, which document the rise to global ecological prominence of green algal phytoplankton (36), as well as the diversification of macroscopic seaweeds (both florideophyte reds and greens (62, 63)) and animals (64-66). The transition from predominantly cyanobacteria to green phytoplankton has been hypothesized to reflect increasing nutrient availability, a conjecture supported by a state change in the abundance of phosphorite deposits during the Ediacaran Period (67). Concomitant increase in oxygen availability, recorded by multiple geochemical signatures (55) fulfills another prediction of increased productivity.

The combination of increasing oxygen and food supply likely facilitated the radiation of animals (34, 68), and metazoan evolution, in turn fed back onto seaweed diversification. Indeed, the Paleozoic evolution of multicellularity and macroscopic growth in ulvophyceans may have been triggered by the introduction of novel grazing pressures by animals in the late Cambrian and Ordovician (30, 66). This may have started an evolutionary arm race between grazers and algae, which would have resulted in increasingly complex feeding and defense strategies, such as larger thalli, larger cells, and mineralized skeletons (49, 59). The relatively late appearance of large grazing animals in the the late Cambrian and Ordovician, such as jawed polychaetes (30, 66) would explain the considerable time lag between the origin of ulvophyceans, and further diversification and origin of crown group ulvophyceans. This scenario is consistent with the predation hypothesis of Stanley (69), who postulated that grazing by benthic animals resulted in a mutual feedback system that drove increased diversity of both macroalgae and animals. The spread of calcium carbonate skeletons in both red and green algae is consistent with the predation hypothesis. Macroscopic growth may also have been facilitated by other factors, including competition for space in benthic habitats, pressure for upright growth to overshadow benthic unicellular algae, a means of increasing nutrient uptake from the water column, or a means of spreading reproductive propagules more widely. A possible mechanism of evolution of attached ulvophycean algae from planktonic ancestors (70) is through intermediary unattached (pleustophytic) macroalgal forms (71), which is consistent with the absence of obvious holdfasts from early ulvophycean fossils (30). In turn, the proliferation of green macroalgae may have resulted in more efficient energy transfer, richer food webs, and the modified shallow marine ecosystems, which may have allowed the evolution of larger and more complex animals (30, 72).

The long-term isolation of ulvophycean lineages in Cryogenian refugia could explain the independent evolution of macroscopic growth in the different clades using radically different mechanisms, as inferred from our phylogeny. Our phylogenetic results, combined with the fossil record indicates that there may have been a time lag of 100 million years or more after the early diversification of ulvophyceans where these microbial ulvophyceans persisted, but never became dominant, followed by transitions to macroscopic growth, which may have occurred in different periods in the different lineages. Our analyses indicate that macroscopic crown group ulvophyceans may have originated between the early Paleozoic (e.g. Bryopsidales) and early Mesozoic (e.g. Cladophorales). This hypothesis is supported by the fossil record, which shows that seaweed morphogroups of the Ediacaran remained largely unchanged during the Cambrian, but that there was a major replacement of this early seaweed flora concomitant with the Great Ordovician Biodiversification Event (30). These observations are also consistent with the fact that the earliest fossils that can be reliably linked to extant ulvophycean clades (Bryopsidales, Dasycladales, and Ulotrichales) are only found from the Ordovician onward (but see discussion on *Proterocladus* above); most (but not all) have calcium carbonate skeletons.

In conclusion, our phylogenetic analysis of the green algae, based on the most comprehensive nuclear gene dataset to data, provides a general picture on the evolution of green seaweeds in which the origin and early diversification of the ulvophyceans likely took place late in the late Tonian and Cryogenian, followed by a marked Ordovician diversification of macroscopic crown group taxa, including those that produce calcium carbonate skeletons and are well preserved in the fossil record. That this trajectory is similar to that of animals reminds us not to forget these organisms when we think about the environmental circumstances of early green algal evolution. Finally, our analyses provide a phylogenetic framework to study the evolution of the genetic toolkit for multicellularity and macroscopic growth in green seaweeds, a group that has been largely neglected in studies of large-scale gene discovery (73).

## Materials and Methods

### Dataset retrieval, RNA extraction and sequencing

DNA sequence data were mined from 15 genomes and 40 transcriptomes, 14 transcriptomes were generated for this study based on cultured strains or freshly collected specimens (*SI Appendix, Material and Methods*), while the remaining data were retrieved from publicly available repositories (*SI Appendix*, Table S1). RNA extractions follow (74). RNA quality and quantity were assessed with Qubit and Nanodrop spectrophotometer, and integrity was assessed with a Bioanalyzer 2100. RNA-seq libraries were sequenced as reported in Table S2.

### Transcriptome assembly, frameshift error correction and ORF detection

At the time of the experiment, pre-assembled transcriptomes were only available for *Acrosiphonia* sp., *Blastophysa rhizopus* and *Caulerpa taxifolia* (Table S1). All remaining assemblies were performed in house, starting from the raw reads using a custom semi-automated pipeline (*SI Appendix, Material and Methods*). For each of the 40 transcriptomes, transcripts were clustered with CD-HIT-EST v. 4.6.1 (75) with a similarity cut-off of 97.5%, and only the longest transcript was retained for downstream analysis as representative of the cluster. After taxonomic profiling of the transcripts (*SI Appendix, Material and Methods*), only eukaryotic transcripts were retained for downstream analysis, bacterial transcripts and transcripts lacking sequence similarity to known proteins were discarded.

Transcripts with putative frameshift errors were identified after initial processing in TRAPID (76), and transcripts carrying a putative frameshift error were corrected with FrameDP 1.2.2 (77), using the *Chlamydomonas* proteome as reference to guide the frameshift correction step (*SI Appendix, Material and Methods*). Each transcript was translated into the corresponding amino acid sequences with the transeq algorithm from the EMBOSS package, using the appropriate translation table, and added to the proteome data of the 15 genomes, resulting in 1,228,821 amino acid sequences.

### Gene Family inference

Sequences were used to build a custom PLAZA 4.0 instance (78), and single-copy families were selected by identifying the 620 picoPLAZA single-copy genes (79) (*SI Appendix, Material and Methods*). To remove potential contaminants in the transcriptomic data from the single-copy gene families, only sequences that were classified as “Viridiplantae” after an additional sequence similarity search with the GhostKOALA (80) webserver were retained for downstream analyses. To further reduce the residual redundancy of the transcriptome datasets in the single-copy gene families, for each gene family the nucleotide sequences of each species were collapsed with CAP3 (81), using stringent parameters to avoid artefactual creation of chimeras: gap penalty 12 (-g) and overlap percent identity cutoff 98% (-p). This set of 620 quasi single-copy genes was used for the downstream phylogenetic analyses.

### Sequence aligments and filtering

Amino acid sequences of the 620 single-copy genes were aligned with MAFFT v. 7.187 (82), using accuracy-oriented parameters (--localpair --maxiterate 1,000) and an offset value (--ep) of 0.075. To identify and trim eventual residual in-paralogs, we followed a phylogeny-guided approach, in which alignments were manually cured to retain only full length or fragments of orthologous single-copy genes (*SI Appendix, Material and Methods*). After filtering, 539 high-confidence single-copy genes out of the 620 initial quasi single-copy genes were retained (hereafter referred to as “coreGF”). The coreGF dataset formed the basis for the construction of seven derived datasets based on different filtering approaches (*SI Appendix, Material and Methods*, Table S3).

### Phylogenetic Analysis

The eight datasets were analyzed using a supermatrix and a coalescence-based phylogenetic approach. ML supermatrix analyses were performed using IQtree with two settings: 1) a gene-wise partitioned analysis (83) was performed, assigning the best substitution model inferred to each partition; 2) an analysis using mixture models was performed using an LG+F+G plus a C20-profile mixture model of substitution rates (84). To estimate the best substitution model of each partition, ML trees were built with IQtree for each single-copy gene, inferring the best model and rate of heterogeneity across sites. All ML analyses were run with 1,000 ultra-fast bootstrap and SH-aLRT branch test replicates.

Gene trees were used also for the coalescent-based analyses, using ASTRAL v. 5.6.1 (85). First, for each ML gene tree, low support branches (ultra fast BS support < 10) were collapsed with Newick Utilities v. 1.6 (86). Branches contracted in the ML gene trees were removed as well from the pool of the corresponding 1,000 bootstrap trees generated during the ML reconstruction. Then, two independent runs were performed either using the ML tree for each gene (BestML), or using the multilocus bootstrap support (MLBS) approach. For the MLBS analysis, 100 replicates were run (-r) starting from the 1,000 contracted bootstrap trees for each gene, allowing gene and site resampling (--gene-resampling flag).

Statistical tests for rejecting the null hypothesis of polytomies were performed in ASTRAL (-t 10 flag), following (87). Briefly, for each of the eight datasets, the coalescence-based MLBS tree was tested (-q flag) by random sampling subsets of ML gene trees representing 1-100% of the total gene families in the dataset, with a minimum of 20 ML gene trees per subset. For each dataset and for each subset, ten independent replicates were generated, and analyzed with ASTRAL on BestML mode with the –t 10 flag –q flag to score the MLBS trees. The support for a number of key relationships was analyzed in each subset of the datasets and the median of the p-values for each subset of trees was calculated.

Different topologies for a number of key relationships were tested using Approximately Unbiased (AU) tests (88) implemented in IQtree, with 100,000 RELL resamplings (89). In addition to the AU test, the gene-wise log-likelihood scores and the percentage of genes supporting alternative topologies were calculated as outlined by (90) and (42).

### Calibrated phylogenetic tree

Due to the high computational cost, the molecular clock analysis was restricted to the 539 coreGF scaffold trim data set. Clock-likeliness of each gene was assessed with the package SortDate (91) against the ML supermatrix and the coalescence-based topologies, scoring the trees on minimal conflict, low root-to-tip variance, and discernible amounts of molecular evolution. The 10 most clock-like genes for each topology were concatenated and subjected to relaxed clock analyses (2,806 and 2,857 amino acid residues for the ML supermatrix and the coalescence-based topology, respectively). Node calibrations were transferred from fossil information and from node age estimates from previous studies (Table S5). All analyses were run with the same set of calibration nodes, except for the UB (*Proterocladus*) and the RT (root age, i.e.: Streptophyta-Chlorophyta split) nodes. The UB node was either constrained (UB_1_) or unconstrained in time (UB_0_). For the root age, three different priors were used (RT_1_-RT_3_), or the root was left unconstrained (RT_0_) (Table S5).

Relaxed molecular clock analyses were run with PhyloBayes 4.1b (92). Two sets of analyses were run on two fixed topologies (the ML supermatrix and the coalescence-based topologies), using the set of clock-like genes for each topology. Both lognormal autocorrelated clock (-ln flag) and uncorrelated gamma multiplier clock (-ugam flag) models were tested for each dataset, and the models were run with either LG+Γ4 or CATGTR+Γ4 models of amino acid substitutions. In total, 64 different analyses were run (Table S6) to test the influence of different models, and of root and key prior ages on the age estimations. For each analysis, two distinct MCMC chains were run for at least 10,000 generations. The convergence of the log likelihoods and parameters estimates were tested in PhyloBayes. Chains were summarized after discarding the first 750 generations as burn-in.

The ultrametric trees were used to guide the ancestral state reconstruction of the ecological and cyto-morphological traits in Phytools (93). The posterior probabilities of the ancestral state of each node were calculated from summaries of 1,000 replicates of simulated stochastic character map (make.simmap), using empirical Bayes method under the ADR model, which permits backward and forward rates between states to have different values.

## Acknowledgements

This work was supported by Ghent University (BOF/01J04813) with infrastructure funded by EMBRC Belgium—FWO project GOH3817N (to O.D.C.), the European Union’s Horizon 2020 research and innovation programme under the Marie Skłodowska-Curie grant agreement No. H2020-MSCA-ITN-2015-675752, the Australian Research Council (DP150100705 to H.V.), and the National Science Foundation (GRAToL 10136495 to C.F.D.). We thank Endymion Cooper for help with generating transcriptomic data. The University of Dundee is a registered Scottish charity, No. 050196. Part of this work has been included and presented in the PhD dissertation of A.D.C. (94).

## Supplementary Information for

### Supplementary text 1: Material and methods

#### Sampling and culture conditions

14 transcriptomes were generated for this study, based on cultured or freshly collected specimens. *Acetabularia acetabulum, Oltmannsiellopsis viridis* and *Scotinosphaera lemnae* cultures were grown at 20 °C and 12-h light/12-h dark cycle in Ace-25 medium (1) and 1.5% agar-solidified Bold Basal medium (2), respectively. *Marsupiomonas* sp., *Pedinomonas minor* and the pedinophyte strain YPF-701 (NIES Microbial Culture Collection strain NIES-2566) were cultured in Guillard’s F/2 medium at 20 °C and 14-h light/10-h dark cycle. For *Ostreobium* sp. HV05042, *Halimeda discoidea* and *Codium fragile* freshly collected material was used for RNA extraction. Collection details are given in (3). Unicellular microscopic algae cultures (*Marsupiomonas* sp., *Oltmannsiellopsis viridis, Pedinomonas minor*, pedinophyte strain YPF-701 and *Scotinosphaera lemnae*) were harvested during their exponential phase. Whole macroscopic seaweed specimens were harvested from cultures (*Acetabularia acetabulum*) or collected in their natural environment (*Codium fragile, Halimeda discoidea* and *Ostreobium* sp. HV05042) and ground in liquid nitrogen for RNA extraction.

#### Transcriptome assembly and taxonomic profiling

Assemblies were performed from the raw reads using a custom semi-automated pipeline, consisted of the following steps. Quality of the raw reads were assessed with FastQC v.0.10.1 (http://www.bioinformatics.babraham.ac.uk, last accessed March 1, 2019). Low-quality reads (average Phreds quality score below 20) and low quality read ends were trimmed with Fastx v.0.0.13 (https://github.com/agordon/fastx_toolkit, last accessed March 1, 2019). Trimmed reads shorter than 30 bp were discarded. Transcriptome *de novo* assembly of *Botryococcus braunii, Chlorokybus atmophyticus, Ulva linza* (Roche 454 data) were performed with CLC Genomics Workbench version 7.5.1 (http://www.clcbio.com, last accessed on March 1, 2019), using a word size of 63 and standard parameters. Transcriptome *de novo* assembly for the remaining species were performed with Trinity 2.1.1 (4), in SE or PE mode were appropriate depending on the RNA-seq library type, after *in silico* reads normalization.

Taxonomic profiling of the transcripts was performed using the following protocol: first, the transcripts were compared to the NCBI non-redundant protein database by sequence similarity searches, using Tera-BLAST DeCypher (Active Motif, USA); then, for each transcript, sequence similarity searches were combined with the NCBI Taxonomy information of the top ten BLAST hits in order to discriminate between eukaryotic and bacterial transcripts or transcripts lacking similarity to known protein-coding genes.

#### Frameshift errors correction and ORF detection

Transcripts with putative frameshift errors were identified after initial processing in TRAPID (5), using the *Chlamydomonas reinhardtii* proteome as reference database for the sequence similarity searches. Transcripts carrying a putative frameshift error were corrected with FrameDP 1.2.2 (6), using the *Chlamydomonas* proteome as reference to guide the frameshift correction step. The longest coding frame was detected with a custom java script plugged into TRAPID using four different translation tables (1, 6, 11 and 16), to take into account alternative nuclear genetic codes (7). Briefly, for each transcript coding frames were predicted with each of the four translation tables. Then, for each transcript, the longest coding sequences detected were retained for the downstream analysis, and the corresponding translation table recorded. Concordance with orthologous sequences predicted by the TRAPID pipeline by sequence similarity searches was also taken into account.

#### Gene Family inference

Sequences were used to build a custom PLAZA 4.0 instance (8) as follows. All-against-all sequence similarity comparison was executed with DIAMOND v. 0.9.18 (9), and the similarity matrix was used to infer homologous relationships among proteins using the graph-based Markov clustering method implemented in Tribe-MCL v. 10-201 (10) with parameters: -scheme 4 –I 2. This resulted in the clustering of 976,181 protein sequences (79.5% of the total proteins) into 69,462 gene families, leaving 252,640 singleton proteins. A procedure was applied to identify and flag outlier proteins from gene families and subfamilies if they showed similarity only to a minority of all family members (11). Single-copy families were selected by identifying the 620 picoPLAZA single-copy genes (12). At this point, to remove potential contaminants in the transcriptomic data from the single-copy gene families, only sequences that were classified as “Viridiplantae” after an additional sequence similarity search with the GhostKOALA (13) webserver were retained for downstream analyses. To further reduce the residual redundancy of the transcriptome datasets in the single-copy gene families, for each gene family the nucleotide sequences of each species were collapsed with CAP3 (14), using stringent parameters to avoid artefactual creation of chimeras: gap penalty 12 (-g) and overlap percent identity cutoff 98% (-p). This set of 620 quasi single-copy genes was used for the downstream phylogenetic analyses.

#### Sequence alignment: identification and trimming of residual in-paralogs

To identify and trim eventual residual in-paralogs, we followed a phylogeny-guided approach. As a first step, a Maximum Likelihood (ML) tree for the resulting alignments was obtained with IQtree v. 1.6.0 (15), inferring the best model and allowing invariable sites and free rate of heterogeneity across sites (16, 17), with parameters: number of ultra-fast bootstrap replicates (-bb) 1,000 (18) and SH-aLRT branch test (19) with 1,000 replicates (-alrt). The resulting trees were visualized in MEGA v. 6 (20) and the corresponding alignments were inspected and processed in Geneious v. 8.0.5 (Biomatters Ltd., https://www.geneious.com/).

Alignments were manually cured to retain only full length or fragments of orthologous single-copy genes: (1) except for *Mesostigma viride* and *Chlorokybus atmophyticus*, which were allowed to cluster with either the Streptophyta or the Chlorophyta, Streptophyta and Chlorophyta monophyly was enforced for the remaining species (e.g. *Arabidopsis thaliana* sequences clustering within the Chlorophyta would be excluded) as obvious sign of paralogy or contamination of environmental samples. (2) For a species with two or more overlapping sequence, one in the expected phylogenetic position (according to orthologous sequences of other species in the same taxon), the other one in a conflicting position, the conflicting sequences were removed. In case of overlapping sequences with concordant phylogenetic signal, regardless of their phylogenetic position, but always enforcing Streptophyta and Chlorophyta monophyly, the longest one was retained, if full-length, otherwise both were retained. In case of conflicting but non-overlapping sequences, both sequences were retained and scaffolded. (3) Gene family alignments with sequences from less than 30 species (ca. half of the total number of species) were discarded. (4) Alignments composed by two or more near-identical paralogs (e.g.: ribosomal subunit proteins) and where confident segregation of the paralogs was difficult were discarded.

#### Sequence alignment: construction of seven derived datasets

The coreGF dataset formed the basis for the construction of seven derived datasets based on different filtering approaches (Table S3): coreGF unscaffold, composed by 539 single-copy genes, with partial sequences removed; coreGF scaffold, composed by 539 single-copy genes, with partial sequences scaffolded; coreGF unscaffold TRIM, as coreGF unscaffold, but with less conserved regions filtered; coreGF scaffold TRIM: as coreGF scaffold, but with less conserved regions filtered; ulvoGF unscaffold, composed by a 355 single-copy genes subset of coreGF focussing on Ulvophyceae, with partial sequences removed; ulvoGF scaffold, composed by a 355 single-copy genes subset of coreGF focussing on Ulvophyceae, with partial sequences scaffolded; ulvoGF unscaffold TRIM, as ulvoGF unscaffold, but with less conserved regions filtered; and ulvoGF scaffold TRIM, as ulvoGF scaffold, but with less conserved regions filtered.

Poorly aligned regions were removed to obtain the corresponding trimmed datasets by trimming the amino acid alignments with TrimAl v. 1.2 (21), with parameters: gapthreshold 0.75 and simthreshold 0.001. Partial sequences from the transcriptomes were either scaffolded (scaffolded dataset) or removed (unscaffolded dataset), resulting in a more comprehensive and a more conservative version of the coreGF and ulvoGF datasets, respectively. Briefly, each gene family with more than one sequence per species (398 gene families), was processed independently for orthology-guided scaffolding with a custom script. First, an all-against-all sequence similarity search was performed with BLASTp v. 2.5.0+ (22), using an e-value cut-off of 10e^-5^. The sequence with the highest bitscore was selected as reference for that gene family. Then, for each species with more than one sequence, the sequences were concatenated according to their relative position to the reference sequence, based on the BLASTp alignments. This resulted in gene families with only one sequence per species. This approach resulted in two types of alignments: a conservative “unscaffold” dataset, where within each gene family, species with more than one sequence were discarded, and a more comprehensive but potentially noisier “scaffold” dataset, where multiple sequences of the same species were scaffolded. Unscaffold and scaffold datasets were re-aligned with MAFFT, using accuracy-oriented parameters (--localpair --maxiterate 1,000 –ep 0.075).

### Supplementary information text 2: Discussion on phylogenetic relationships within the core Chlorophyta

#### Phylogenetic positions of Pedinophyceae and Chlorodendrophyceae

Our phylogenetic analyses recovered the Chlorodendrophyceae and Pedinophyceae as the two earliest diverging lineages of the core Chlorophyta, consistent with some previous phylogenetic analyses based on nuclear- and plastid-encoded rDNA sequences (23) and chloroplast phylogenomic analyses (3, 24, 25). However, the exact phylogenetic position (Chlorodendrophyceae first, or Pedinophyceae first) differed between analyses (supplementary figures). The phylogenetic position of these two lineages, relative to the remaining core chlorophytan lineages is relevant in light of evolution of cell division. In contrast to the prasinophytes where cell division occurs by furrowing, three classes of the core Chlorophyta, including the Chlorophyceae, Trebouxiophyceae and Chlorodendrophyceae, are characterized by a new mode of cell division that is mediated by a phycoplast, which is an array of microtubules oriented parallel to the plane of cell division, determining the formation of a new cell wall (26, 27). A phycoplast is absent in the Pedinophyceae and Ulvophyceae (Trentepohliales have a phragmoplast-like cytokinesis, similar to that found in some Streptophyta). Since our phylogenetic analyses are inconclusive with respect to the earliest diverging lineage of the core Chlorophyta (Pedinophyceae or Chlorodendrophyceae) it remains uncertain whether the phycoplast evolved at the base of the core Chlorophyta and was subsequently lost in the Pedinophyceae, or evolved after the divergence of the Pedinophyceae.

#### Phylogenetic relationships between the Ulvophyceae, Trebouxiophyceae and Chlorophyceae (UTC)

The Trebouxiophyceae and Chlorophyceae were unambiguously recovered as monophyletic groups, with the Trebouxiophyceae sister to a clade containing the Chlorophyceae and Ulvophyceae. Previous phylogenetic studies have not been able to resolve these relationships unambiguously (27). Ultrastructural evidence has been interpreted as either providing support for a sister relation between Chlorophyceae and Trebouxiophyceae based on the shared presence of a phycoplast and a non-persistent mitotic spindle (26), or between Trebouxiophyceae and Ulvophyceae based on a counterclockwise orientation of the flagellar basal bodies (28). Phylogenies based on nuclear rDNA sequences have been inconclusive, with the branching order of the UTC classes depending on taxon sampling, alignment methods and phylogenetic methods. Some early 18S rDNA phylogenies showed a sister relationship between Chlorophyceae and Trebouxiophyceae (e.g., 29) while others revealed a sister relation between the Chlorophyceae and Ulvophyceae (e.g., 30). A 10-gene (2 plastid and 8 nuclear genes) phylogeny (31) showed a sister relationship between the Chlorophyceae and Ulvophyceae, indicating that the ancestor of the UTC clade had a phycoplast and a non-persistent mitotic spindle, and that both characters have been lost in the Ulvophyceae. The ancestral status of a phycoplast and a non-persistent mitotic spindle is congruent with the presence of these features in the early diverging Chlorodendrophyceae (26). Likewise the ancestor of the UTC clade probably had a counterclockwise orientation of the flagellar basal bodies, evolving to a direct opposite or clockwise orientation in the Chlorophyceae. Again this interpretation is congruent with the presence of a counterclockwise basal body orientation in the Chlorodendrophyceae. Chloroplast phylogenomic analyses have not been able to resolve the branching order of the UTC classes conclusively, with the main uncertainties including monophyly of the Trebouxiophyceae and Ulvophyceae (see below), and the sister clade of the Chlorophyceae (25, 32).

#### Monophyly of the Ulvophyceae and phylogenetic position of the Bryopsidales

The distinct phylogenetic position of the Bryopsidales in our analyses is unexpected from a morphological view-point. The order has long been allied with the Dasycladales based on the shared siphonous structure, and this relationship has been confirmed by small nuclear gene datasets (30, 31, 33), while in chloroplast phylogenomic studies, the position of the Bryopsidales has been unstable (32, 34, 35). Conversely, ultrastructural features of the flagellar apparatus suggested a close relationship between the Dasycladales and Cladophorales. The two orders share a flattened flagellar apparatus, have similar striations on the distal fibers connecting the basal bodies, and lack terminal caps on the anterior surface of the basal bodies (36). In assuming a sister relationship between the Bryopsidales and Dasycladales based on analysis of a 10-gene dataset, Cocquyt et al. (31) hypothesised that the evolution of siphonous growth would have involved two steps, including the development of an enlarged ancestral cell with a macronucleus and cytoplasmic streaming, allowing increased transcription from a single nucleus and distribution of transcripts across the cell, followed by the evolution of multinucleate siphons, which possibly occurred independently in the Bryopsidales and Dasycladales in association with the evolution of complex, macroscopic plants that have outgrown the potential of a single macronucleus. Our phylogenetic results point towards an entirely independent evolution of siphonous organization in Bryopsidales and Dasycladales from an ancestral unicellular green alga. The distinct phylogenetic position of the Bryopsidales, separate from the Trentepohliales, Cladophorales, and Dasycladales is supported by independent molecular data. The latter orders, along with the Scotinosphaerales and *Blastophysa*, are characterized by a non-canonical nuclear genetic code while the Bryopsidales and all other green algae have the canonical code (7, 37), pointing towards a single origin of a non-canonical nuclear code in the green algae. The Bryopsidales and Dasycladales also present fundamental differences in life cycle and cytology, for example in their approaches towards cytoplasmic streaming (microtubules and actin versus actin-only-powered, respectively) (38-43), supporting a more distant relationship between the two orders.

#### Phylogenetic position of the microscopic ulvophycean orders: Ignatiales, Oltmansiellopsidales and Scotinosphaerales

The phylogenetic affinities of the microscopic and mostly unicellular ulvophycean orders Ignatiales (including non-motile unicellular or loose colonial algae living on soil or freshwater macrophytes), Oltmansiellopsidales (including motile and non-motile unicellular or colonial algae from marine or freshwater habitats), and Scotinosphaerales (including non-motile unicellular algae from freshwater or damp terrestrial habitats) are important to understand the evolution of macroscopic growth and multicellularity in the Ulvophyceae. Species in these orders were originally described in the Chlorophyceae or Trebouxiophyceae based on morphological grounds, but molecular phylogenetic studies firmly placed them in the Ulvophyceae (30, 31, 44, 45). Our analyses are consistent with chloroplast phylogenomic analyses in placing the Ignatiales and Oltmansiellopsidales in a clade together with the Ulvales-Ulotrichales (32), but in contrast to chloroplast-based analyses, the Oltmansiellopsidales was found to branch off first in our analyses. In contrast to a two-gene analysis, which recovered the Scotinosphaerales sister to a clade including the Ignatiales, Oltmansiellopsidales, and Ulvales-Ulotrichales (45), our analyses indicate that the Scotinosphaerales are sister to the Dasycladales. Although taxon sampling and the exact phylogenetic positions of microscopic ulvophycean lineages differed from the 10-gene analysis of Cocquyt et al. (31), our results support the hypothesis that macroscopic growth evolved independently in various lineages of ulvophytes from ancestral unicellular green algae by different mechanisms: by developing multicellularity with coupled mitosis and cytokinesis (independently in the Ulvales-Ulotrichales and Trentepohliales), by developing multicellularity with uncoupled mitosis and cytokinesis (siphonocladous organisation in the Cladophorales and *Blastophysa*) or by developing a siphonous architecture (independently in the Bryopsidales and Dasycladales).

### Supplementary tables

**Table S1.**
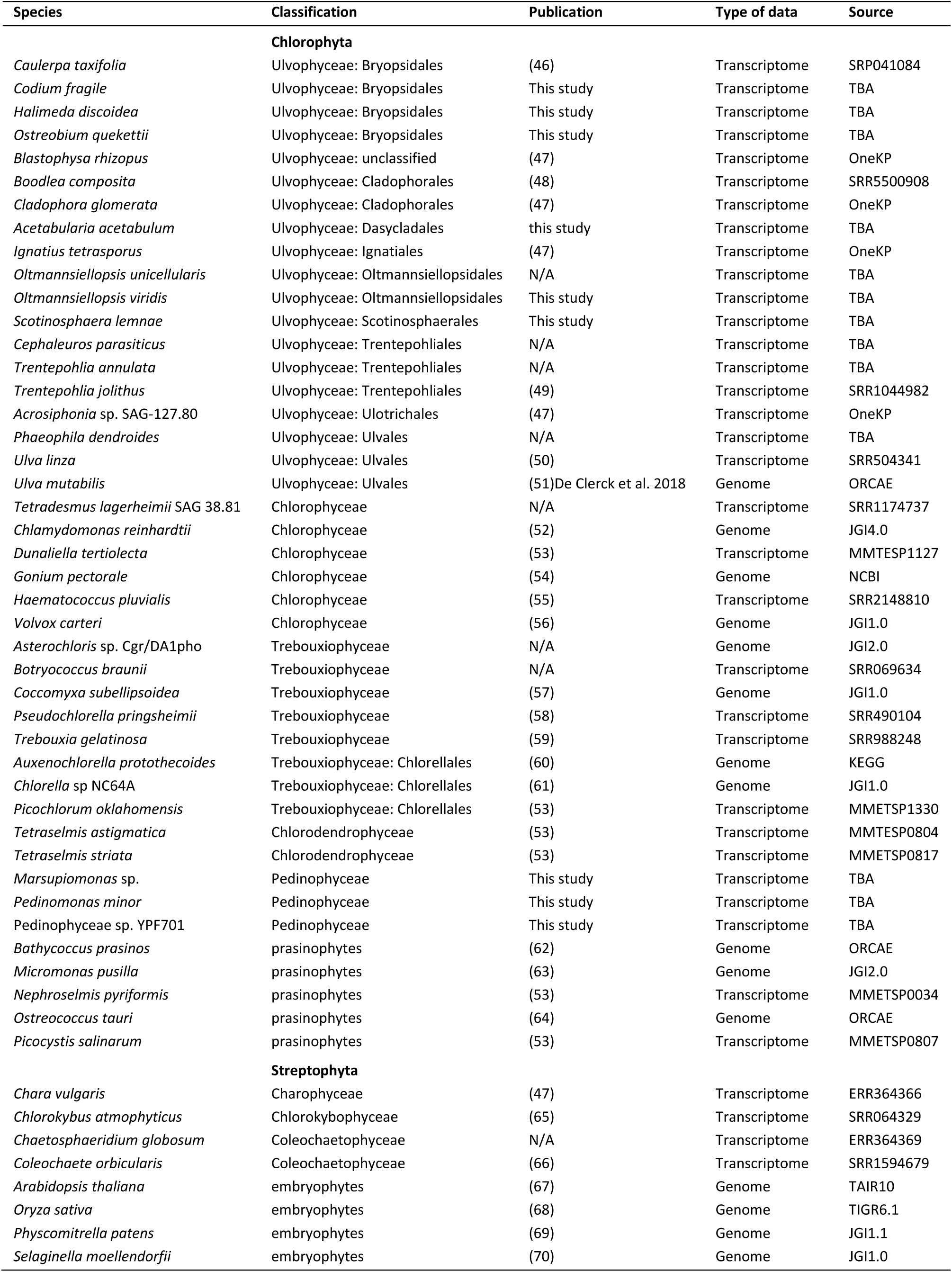
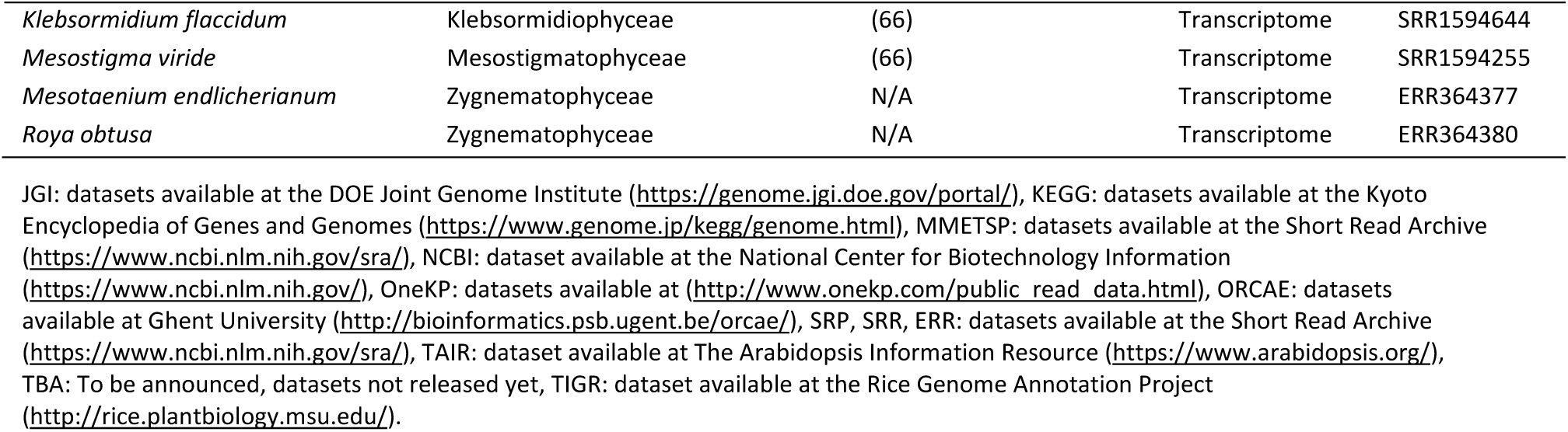
Transcriptomic and genomic datasets used in this study.

**Table S2.**
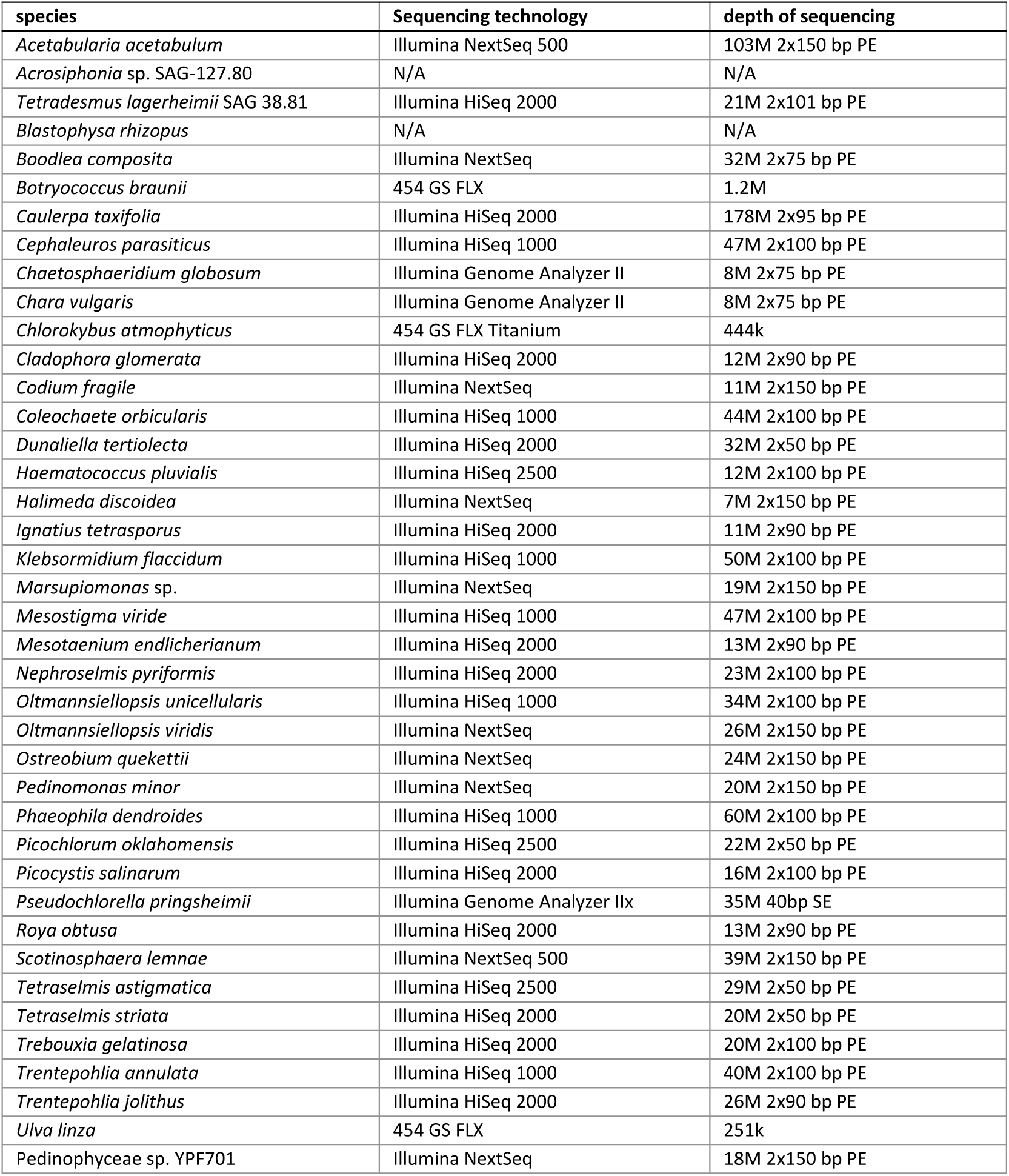
Sequencing technology and read depth of the transcriptome data.

**Table S3.** Alignments metrics of the eight datasets analyzed.

|  | coreGF |  |  |  | ulvoGF |  |  |  |
| --- | --- | --- | --- | --- | --- | --- | --- | --- |
|  | unscaffold |  | scaffold |  | unscaffold |  | Scaffold |  |
|  | untrimmed | trimmed | untrimmed | trimmed | untrimmed | trimmed | untrimmed | Trimmed |
| # genes | 539 | 539 | 539 | 539 | 355 | 355 | 355 | 355 |
| # sites (nt) | 1,751,925 | 342, 501 | 1,877,544 | 342,501 | 1,143,789 | 252,612 | 1,244,520 | 252,612 |
| # sites (aa) | 583,975 | 114,167 | 625,848 | 114,167 | 381,263 | 84,204 | 414,840 | 84,204 |
| % missing data | 65.6 | 27.3 | 69.7 | 19.9 | 43.7 | 24.0 | 54.8 | 15.9 |

nt = nucleotides, aa = amino acid

**Table S4.**
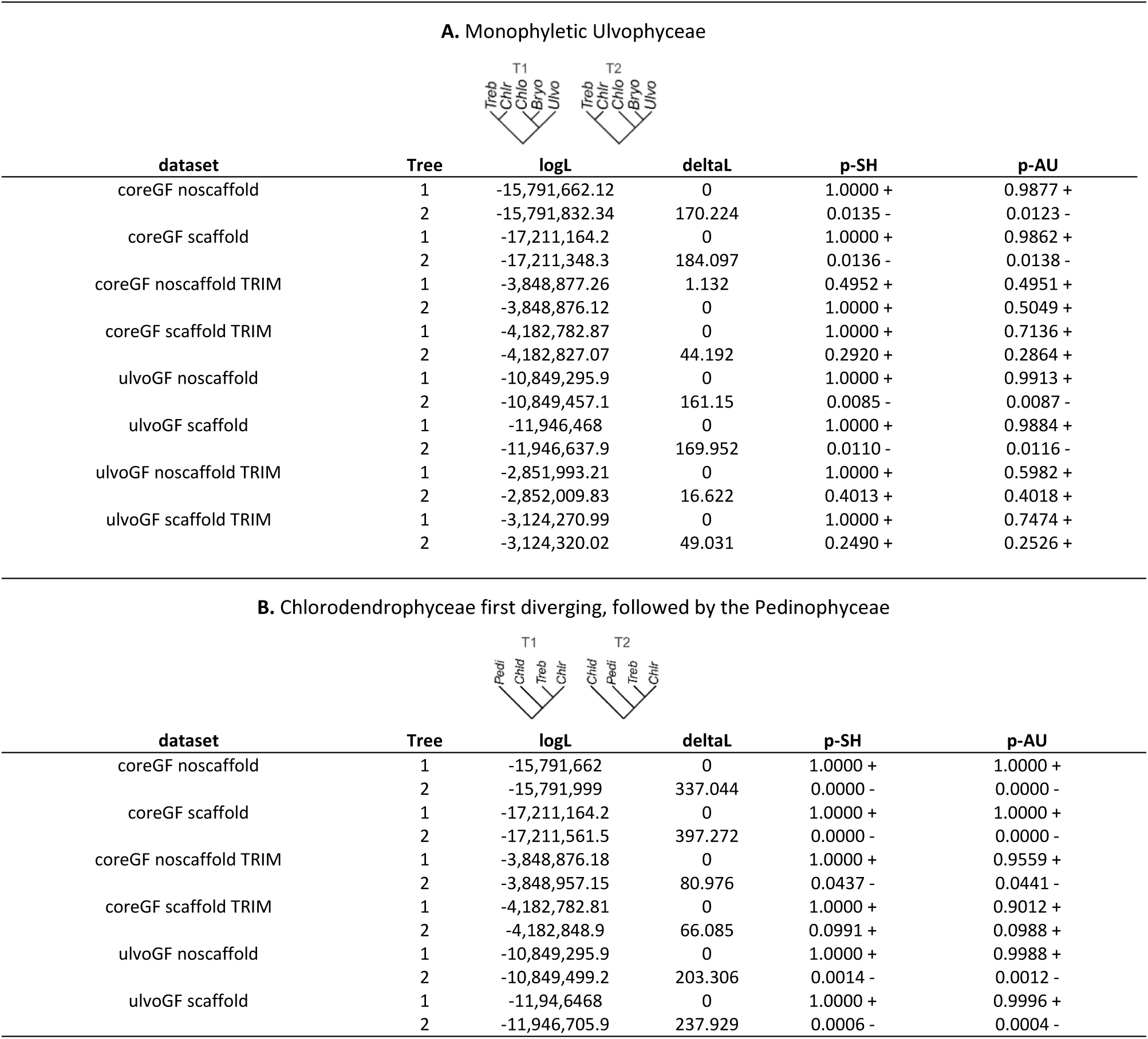
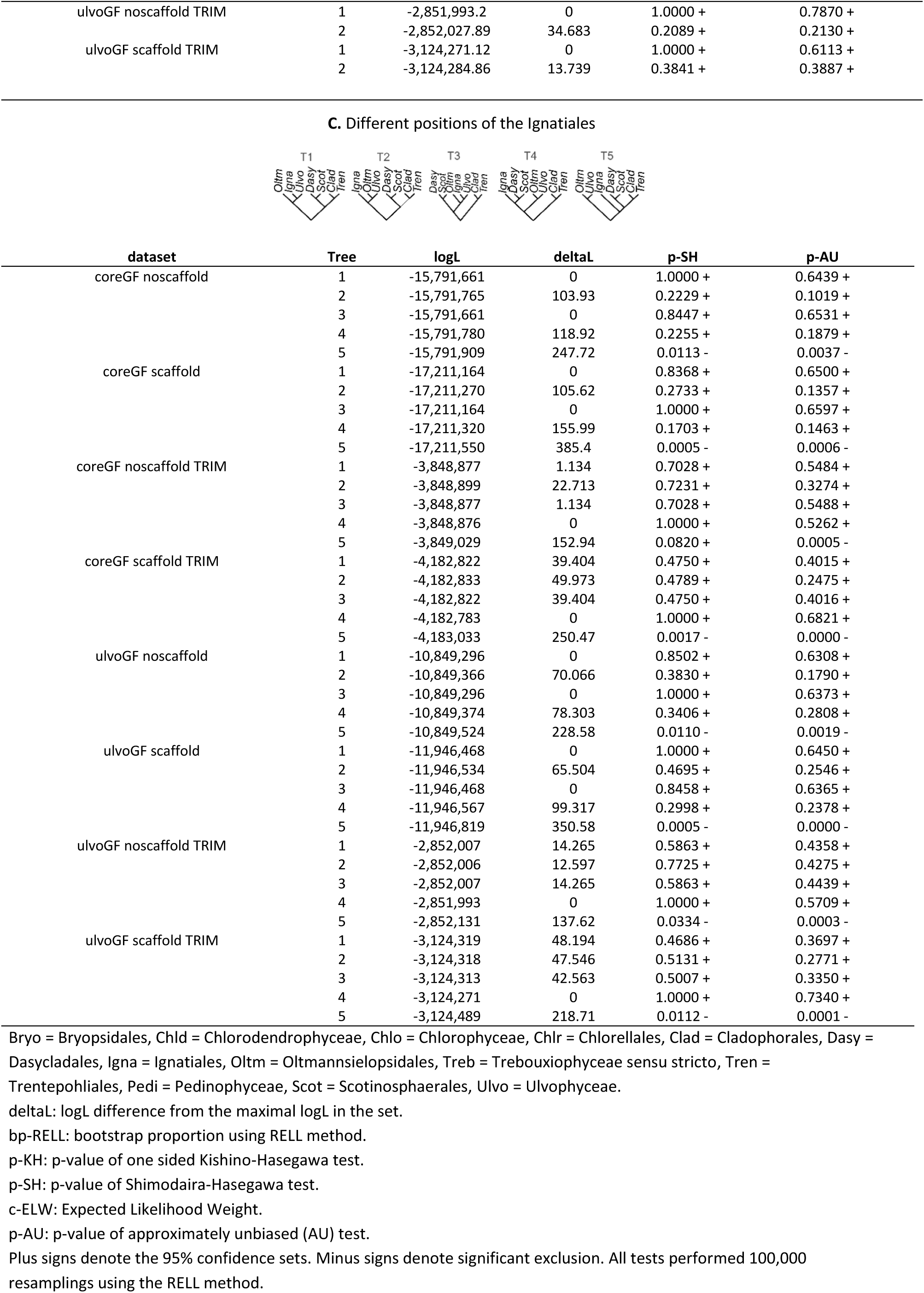
Comparing alternative hypotheses using the Shimodaira-Hasegawa (SH) Test and Approximately Unbiased (AU) Test. Constraint trees include: (A) Monophyletic Ulvophyceae (coalescence-based topology), (B) Chlorodendrophyceae diverging first, followed by the Pedinophyceae, (C) Different positions of the Ignatiales. The topological schemas (hypotheses) are depicted. T1 represents the unconstrained tree topology.

| A. Monophyletic Ulvophyceae |  |  |  |  |  |
| --- | --- | --- | --- | --- | --- |
| dataset | Tree | logL | deltal | p-SH | p-AU |
| coreGF noscaffold | 1 | -15,791,662.12 | 0 | 1.0000 + | 0.9877 + |
|  | 2 | -15,791,832.34 | 170.224 | 0.0135 - | 0.0123 - |
| coreGF scaffold | 1 | -17,211,164.2 | 0 | 1.0000 + | 0.9862 + |
|  | 2 | -17,211,348.3 | 184.097 | 0.0136 - | 0.0138 - |
| coreGF noscaffold TRIM | 1 | -3,848,877.26 | 1.132 | 0.4952 + | 0.4951 + |
|  | 2 | -3,848,876.12 | 0 | 1.0000 + | 0.5049 + |
| coreGF scaffold TRIM | 1 | -4,182,782.87 | 0 | 1.0000 + | 0.7136 + |
|  | 2 | -4,182,827.07 | 44.192 | 0.2920 + | 0.2864 + |
| ulvoGF noscaffold | 1 | -10,849,295.9 | 0 | 1.0000 + | 0.9913 + |
|  | 2 | -10,849,457.1 | 161.15 | 0.0085 - | 0.0087 - |
| ulvoGF scaffold | 1 | -11,946,468 | 0 | 1.0000 + | 0.9884 + |
|  | 2 | -11,946,637.9 | 169.952 | 0.0110 - | 0.0116 - |
| ulvoGF noscaffold TRIM | 1 | -2,851,993.21 | 0 | 1.0000 + | 0.5982 + |
|  | 2 | -2,852,009.83 | 16.622 | 0.4013 + | 0.4018 + |
| ulvoGF scaffold TRIM | 1 | -3,124,270.99 | 0 | 1.0000 + | 0.7474 + |
|  | 2 | -3,124,320.02 | 49.031 | 0.2490 + | 0.2526 + |

| B. Chlorodendrophyceae first diverging, followed by the Pedinophyceae |  |  |  |  |  |
| --- | --- | --- | --- | --- | --- |
| dataset | Tree | logL | deltal | p-SH | p-AU |
| coreGF noscaffold | 1 | -15,791,662 | 0 | 1.0000 + | 1.0000 + |
|  | 2 | -15,791,999 | 337.044 | 0.0000 - | 0.0000 - |
| coreGF scaffold | 1 | -17,211,164.2 | 0 | 1.0000 + | 1.0000 + |
|  | 2 | -17,211,561.5 | 397.272 | 0.0000 - | 0.0000 - |
| coreGF noscaffold TRIM | 1 | -3,848,876.18 | 0 | 1.0000 + | 0.9559 + |
|  | 2 | -3,848,957.15 | 80.976 | 0.0437 - | 0.0441 - |
| coreGF scaffold TRIM | 1 | -4,182,782.81 | 0 | 1.0000 + | 0.9012 + |
|  | 2 | -4,182,848.9 | 66.085 | 0.0991 + | 0.0988 + |
| ulvoGF noscaffold | 1 | -10,849,295.9 | 0 | 1.0000 + | 0.9988 + |
|  | 2 | -10,849,499.2 | 203.306 | 0.0014 - | 0.0012 - |
| ulvoGF scaffold | 1 | -11,94,6468 | 0 | 1.0000 + | 0.9996 + |
|  | 2 | -11,946,705.9 | 237.929 | 0.0006 - | 0.0004 - |
| ulvoGF noscaffold TRIM | 1 | -2,851,993.2 | 0 | 1.0000 + | 0.7870 + |
|  | 2 | -2,852,027.89 | 34.683 | 0.2089 + | 0.2130 + |
| ulvoGF scaffold TRIM | 1 | -3,124,271.12 | 0 | 1.0000 + | 0.6113 + |
|  | 2 | -3,124,284.86 | 13.739 | 0.3841 + | 0.3887 + |

| dataset | Tree | logL | deltaL | p-SH | p-AU |
| --- | --- | --- | --- | --- | --- |
| coreGF noscaffold | 1 | -15,791,661 | 0 | 1.0000 + | 0.6439 + |
|  | 2 | -15,791,765 | 103.93 | 0.2229 + | 0.1019 + |
|  | 3 | -15,791,661 | 0 | 0.8447 + | 0.6531 + |
|  | 4 | -15,791,780 | 118.92 | 0.2255 + | 0.1879 + |
|  | 5 | -15,791,909 | 247.72 | 0.0113 - | 0.0037 - |
| coreGF scaffold | 1 | -17,211,164 | 0 | 0.8368 + | 0.6500 + |
|  | 2 | -17,211,270 | 105.62 | 0.2733 + | 0.1357 + |
|  | 3 | -17,211,164 | 0 | 1.0000 + | 0.6597 + |
|  | 4 | -17,211,320 | 155.99 | 0.1703 + | 0.1463 + |
|  | 5 | -17,211,550 | 385.4 | 0.0005 - | 0.0006 - |
| coreGF noscaffold TRIM | 1 | -3,848,877 | 1.134 | 0.7028 + | 0.5484 + |
|  | 2 | -3,848,899 | 22.713 | 0.7231 + | 0.3274 + |
|  | 3 | -3,848,877 | 1.134 | 0.7028 + | 0.5488 + |
|  | 4 | -3,848,876 | 0 | 1.0000 + | 0.5262 + |
|  | 5 | -3,849,029 | 152.94 | 0.0820 + | 0.0005 - |
| coreGF scaffold TRIM | 1 | -4,182,822 | 39.404 | 0.4750 + | 0.4015 + |
|  | 2 | -4,182,833 | 49.973 | 0.4789 + | 0.2475 + |
|  | 3 | -4,182,822 | 39.404 | 0.4750 + | 0.4016 + |
|  | 4 | -4,182,783 | 0 | 1.0000 + | 0.6821 + |
|  | 5 | -4,183,033 | 250.47 | 0.0017 - | 0.0000 - |
| ulvoGF noscaffold | 1 | -10,849,296 | 0 | 0.8502 + | 0.6308 + |
|  | 2 | -10,849,366 | 70.066 | 0.3830 + | 0.1790 + |
|  | 3 | -10,849,296 | 0 | 1.0000 + | 0.6373 + |
|  | 4 | -10,849,374 | 78.303 | 0.3406 + | 0.2808 + |
|  | 5 | -10,849,524 | 228.58 | 0.0110 - | 0.0019 - |
| ulvoGF scaffold | 1 | -11,946,468 | 0 | 1.0000 + | 0.6450 + |
|  | 2 | -11,946,534 | 65.504 | 0.4695 + | 0.2546 + |
|  | 3 | -11,946,468 | 0 | 0.8458 + | 0.6365 + |
|  | 4 | -11,946,567 | 99.317 | 0.2998 + | 0.2378 + |
|  | 5 | -11,946,819 | 350.58 | 0.0005 - | 0.0000 - |
| ulvoGF noscaffold TRIM | 1 | -2,852,007 | 14.265 | 0.5863 + | 0.4358 + |
|  | 2 | -2,852,006 | 12.597 | 0.7725 + | 0.4275 + |
|  | 3 | -2,852,007 | 14.265 | 0.5863 + | 0.4439 + |
|  | 4 | -2,851,993 | 0 | 1.0000 + | 0.5709 + |
|  | 5 | -2,852,131 | 137.62 | 0.0334 - | 0.0003 - |
| ulvoGF scaffold TRIM | 1 | -3,124,319 | 48.194 | 0.4686 + | 0.3697 + |
|  | 2 | -3,124,318 | 47.546 | 0.5131 + | 0.2771 + |
|  | 3 | -3,124,313 | 42.563 | 0.5007 + | 0.3350 + |
|  | 4 | -3,124,271 | 0 | 1.0000 + | 0.7340 + |
|  | 5 | -3,124,489 | 218.71 | 0.0112 - | 0.0001 - |

Bryo = Bryopsidales, Chld = Chlorodendrophyceae, Chlo = Chlorophyceae, Chlr = Chlorellales, Clad = Cladophorales, Dasy = Dasycladales, Igna = Ignatiales, Oltm = Oltmannsielopsidales, Treb = Trebouxiophyceae sensu stricto, Tren = Trentepohliales, Pedi = Pedinophyceae, Scot = Scotinosphaerales, Ulvo = Ulvophyceae.
deltaL: logL difference from the maximal logL in the set.
bp-RELL: bootstrap proportion using REll method.
p-KH: p-value of one sided Kishino-Hasegawa test.
p-SH: p-value of Shimodaira-Hasegawa test.
c-ELW: Expected Likelihood Weight.
p-AU: p-value of approximately unbiased (AU) test.
Plus signs denote the 95% confidence sets. Minus signs denote significant exclusion. All tests performed 100,000 resamplings using the REll method.

**Table S5.** Node calibrations used for relaxed molecular clock analysis

| Node | Node | calibration | Fossil or transfer | period | prior | Reference |
| --- | --- | --- | --- | --- | --- | --- |
| C | Characeae | C<sub>1</sub> | Transfer from other study | Devonian | U[416-∞[ | (71) |
| L | Land plants (embryophytes) | L<sub>1</sub> | Cryptospore assemblage from the Middle Ordovician (Dapingian) | Ordovician | U[475-∞[ | (71) |
| V | Vascular plants | V<sub>1</sub> | *Baragwanathia longifolia* | Silurian | U[421-∞[ | (71) |
| T<sub>1</sub> | *Botryococcus* stem node | T<sub>1</sub> | *Botryococcus* | Carboniferous | U[299-∞ [ | (72) |
| UA | *Ostreobium* stem node | UA<sub>1</sub> | Transfer from other study | n/a | U[533-425] | (73) |
| UB | Ulvophyceae/Chlorophyceae stem node | UB<sub>1</sub> | *Proterocladus* | Neoproterozoic | U[716-∞ [ | (74) |
|  |  | UB<sub>0</sub> | Absence of calibration | n/a | n/a | n/a |
| RT | Streptophyta-Chlorophyta split | RT<sub>1</sub> | Transfer from other study | n/a | U[1279-1159] | (75) |
|  |  | RT<sub>2</sub> | Transfer from other study | n/a | U[1015-863] | (76) |
|  |  | RT<sub>3</sub> | RT<sub>1</sub>/RT<sub>2</sub> hybrid | n/a | U[1279-863] | n/a |
|  |  | RT<sub>0</sub> | n/a | n/a | n/a | n/a |

**Table S6.**
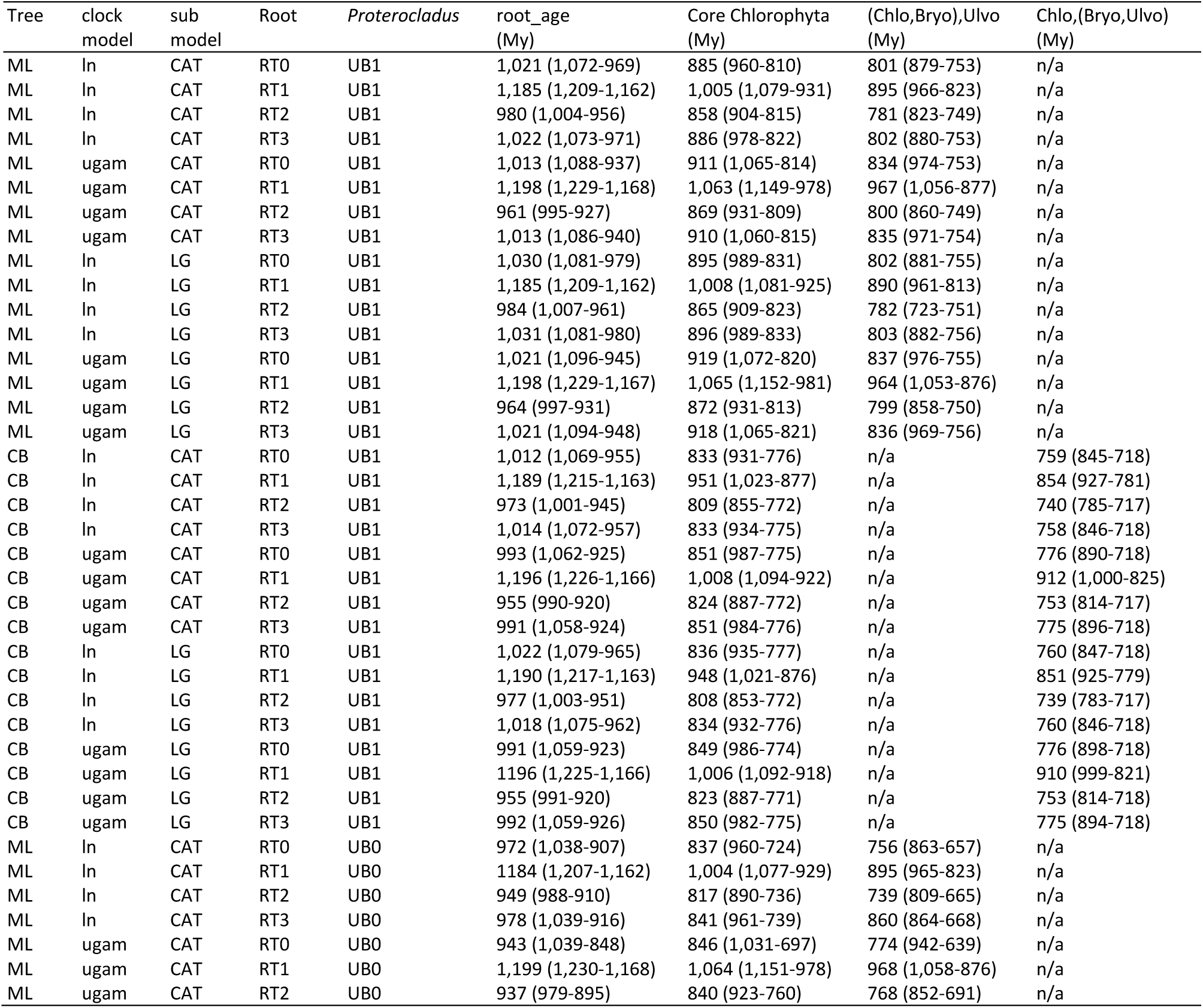
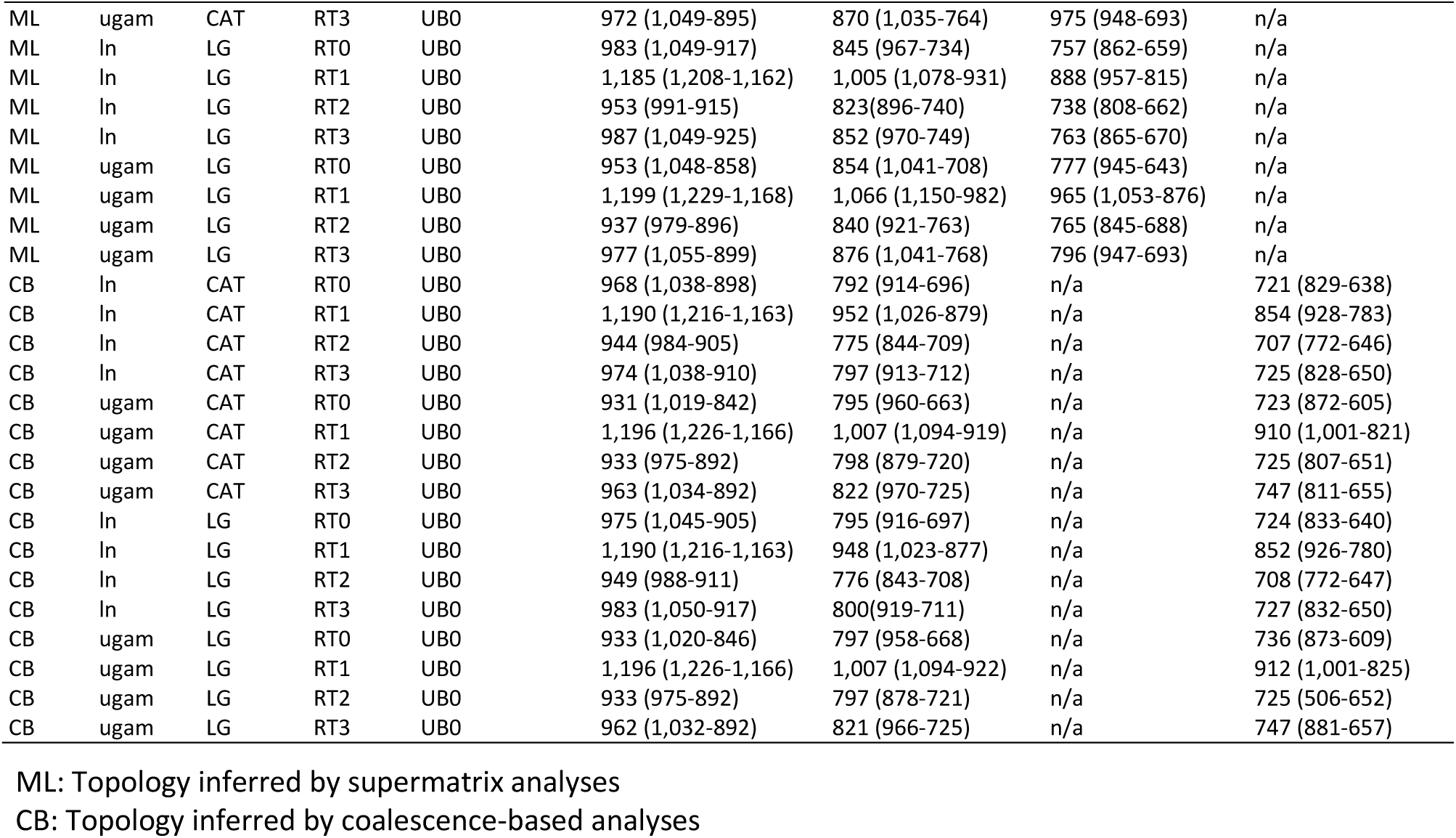
Relaxed molecular clock analyses results.

### Supplementary figures

**Figure S1.**
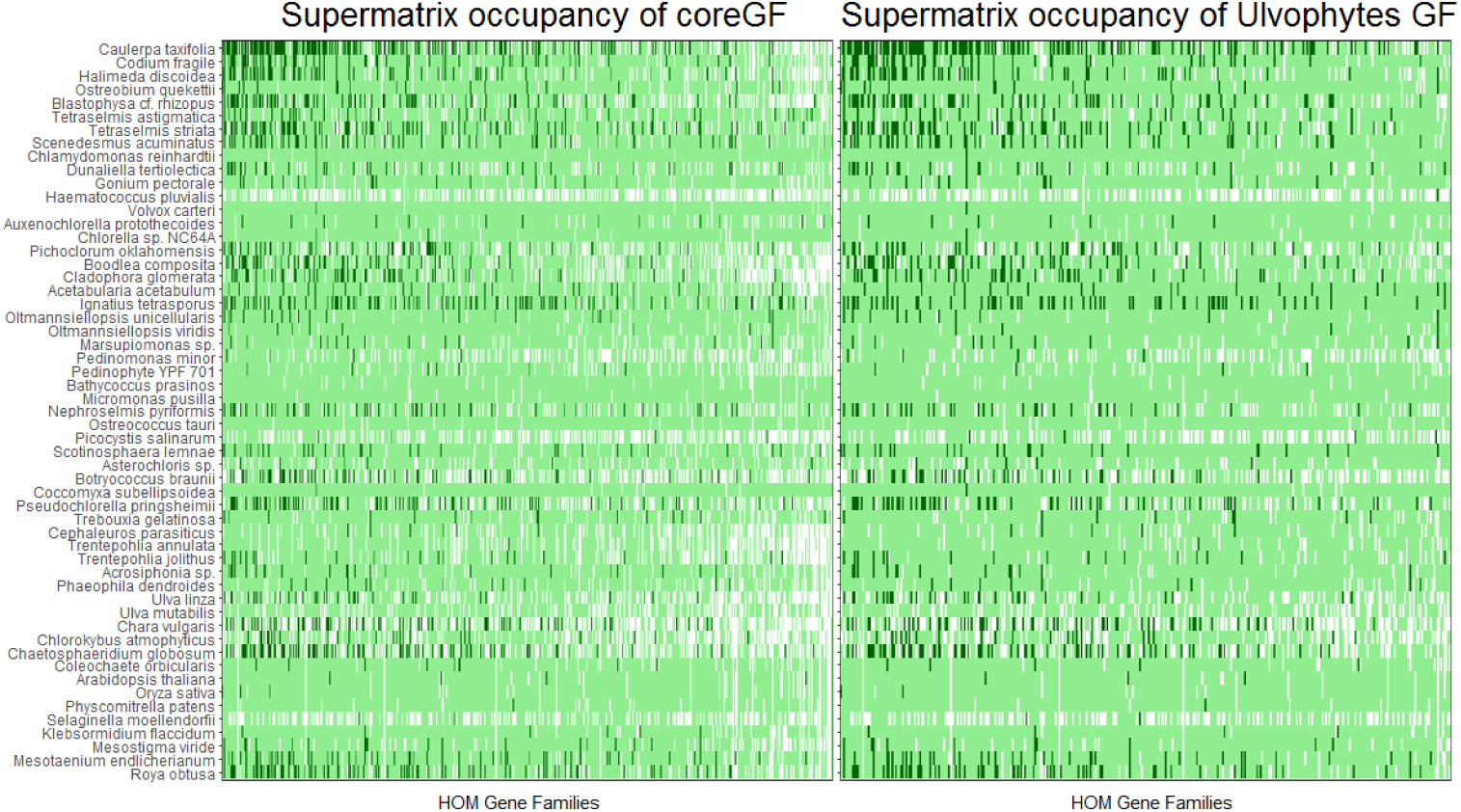
Supermatrix occupancy of the coreGF and ulvoGF datasets. Supermatrix occupancy of the 539 coreGF and 355 ulvoGF single-copy genes. From the left to the right, the genes with highest to lowest taxon occupancy. Dark green: scaffolded sequence (absent from the unscaffold datasets); light green: available sequence; white: missing sequence.

**Figure S2.**
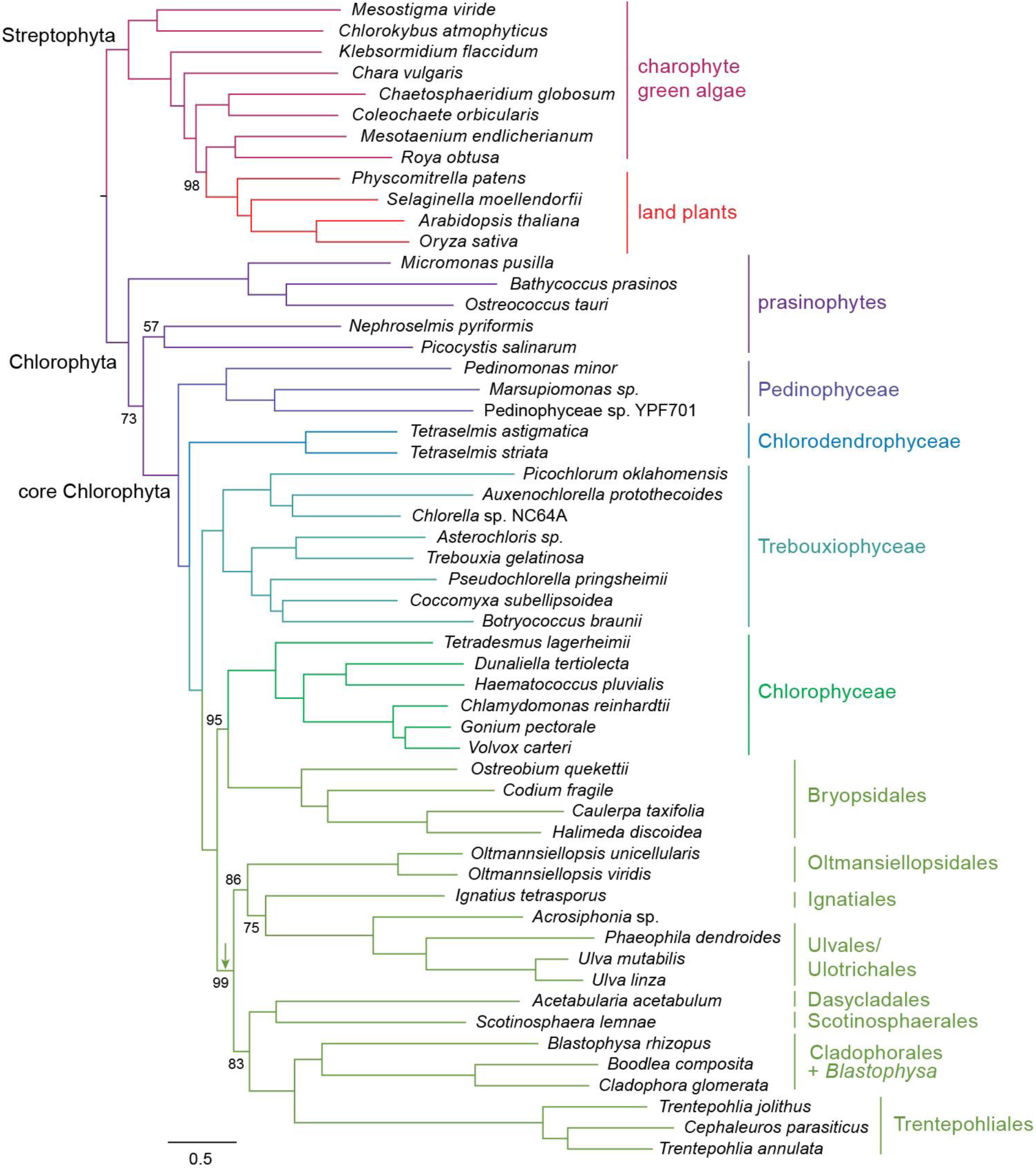
Maximum likelihood phylogenetic tree of the green algae inferred from a concatenated amino acid alignment of 539 nuclear genes (supermatrix analysis of the coreGF-scaffolded-untrimmed data set). Bootstrap support values are shown at the nodes (100% values are omitted). Scale bar indicates estimated number of substitutions per amino acid position. The green arrow indicates the phylogenetic position of the Bryopsidales in the coalescence-based analyses.

**Figure S3.**
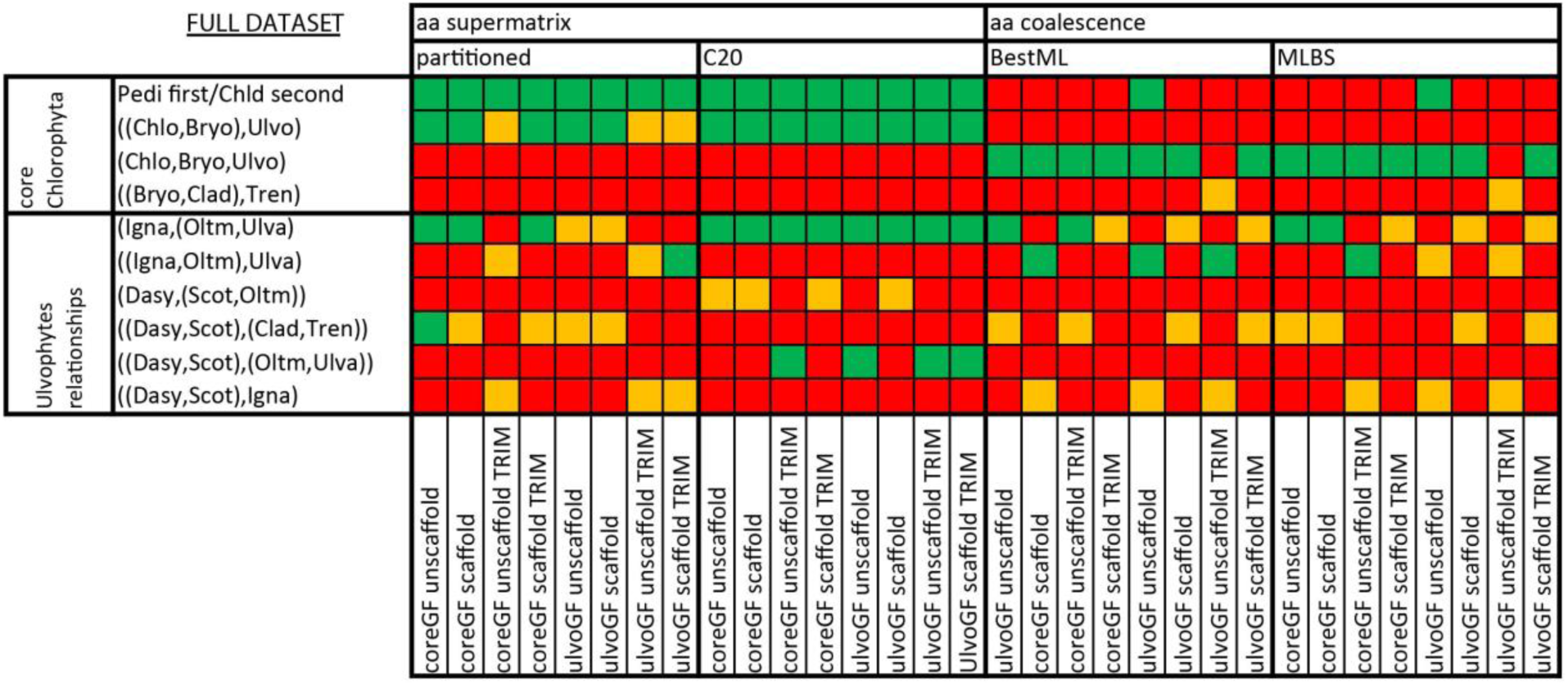
Summary of support for hypotheses of specific chlorophytan relationships, based on supermatrix or coalescence-based analyses of the eight amino acid alignments (Table S3). Relationships are summarized in newick tree format. Green tiles indicate high support (bootstrap values and posterior probabilities > 75% and 0.75, respectively), yellow tiles indicate weak support, while red tiles indicate that the phylogenetic relationship was rejected. Bryo: Bryopsidales; Chld: Chlorodendrophyceae; Chlo: Chlorophyceae; Chlr: Chlorellales; Clad: Cladophorales+*Blastophysa*; Dasy: Dasycladales; Igna: Ignatiales; Oltm: Oltmannsiellopsidales; Pedi: Pedinophyceae; Scot: Scotinosphaerales; Tren: Trentepohliales; Ulva: Ulvales-Ulotrichales; Ulvo: Ulvophyceae excluding Bryopsidales.

**Figure S4.**
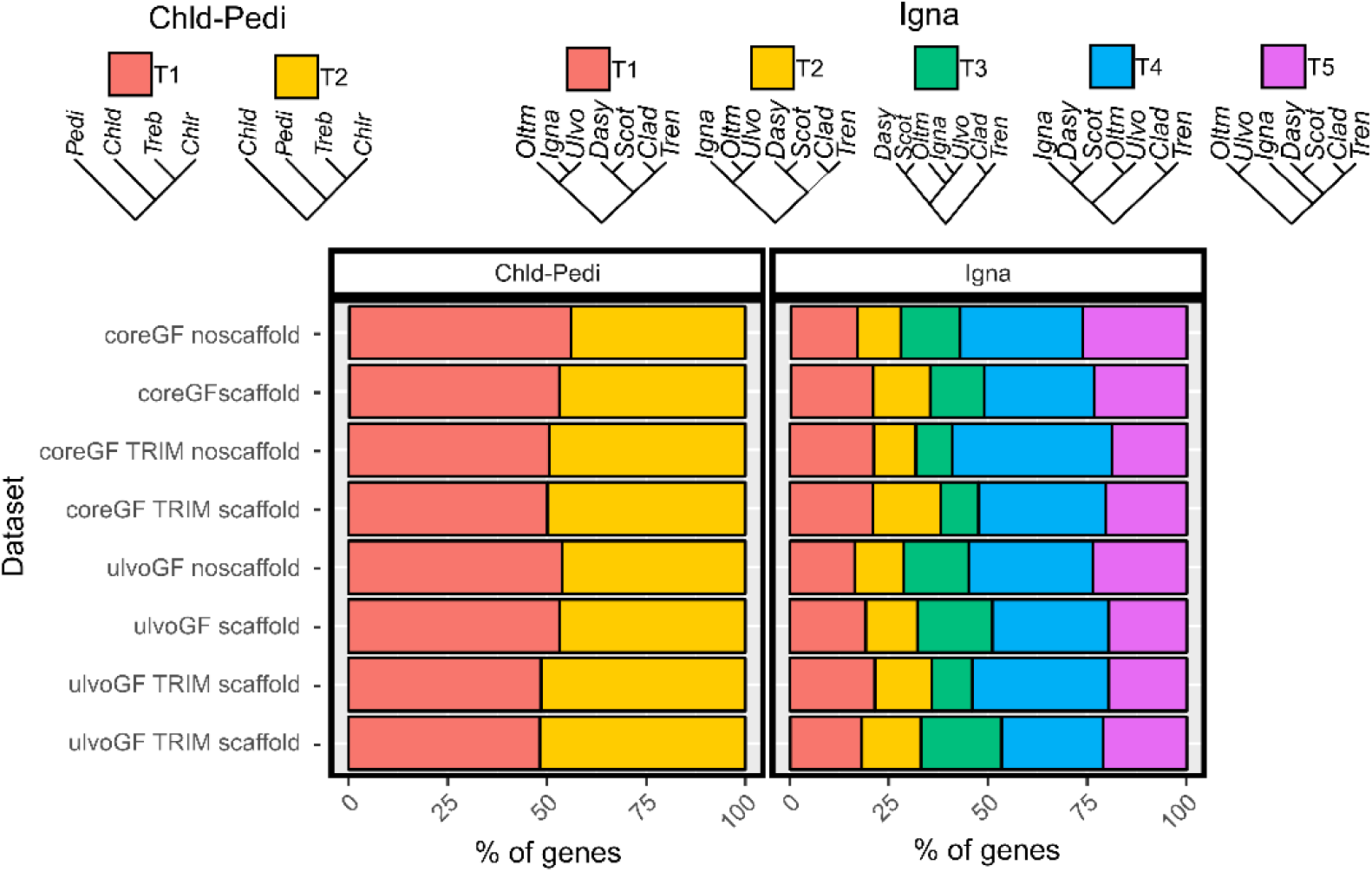
Proportion of genes supporting each of the alternative topologies in pairwise log-likelihood score comparisons. Alternative topologies were also tested using AU-tests (Table S4). Bryo: Bryopsidales; Chld: Chlorodendrophyceae; Chlo: Chlorophyceae; Chlr: Chlorellales; Clad: Cladophorales+*Blastophysa*; Dasy: Dasycladales; Igna: Ignatiales; Oltm: Oltmannsiellopsidales; Pedi: Pedinophyceae; Scot: Scotinosphaerales; Tren: Trentepohliales; Ulvo: Ulvales-Ulotrichales.

**Figure S5.**
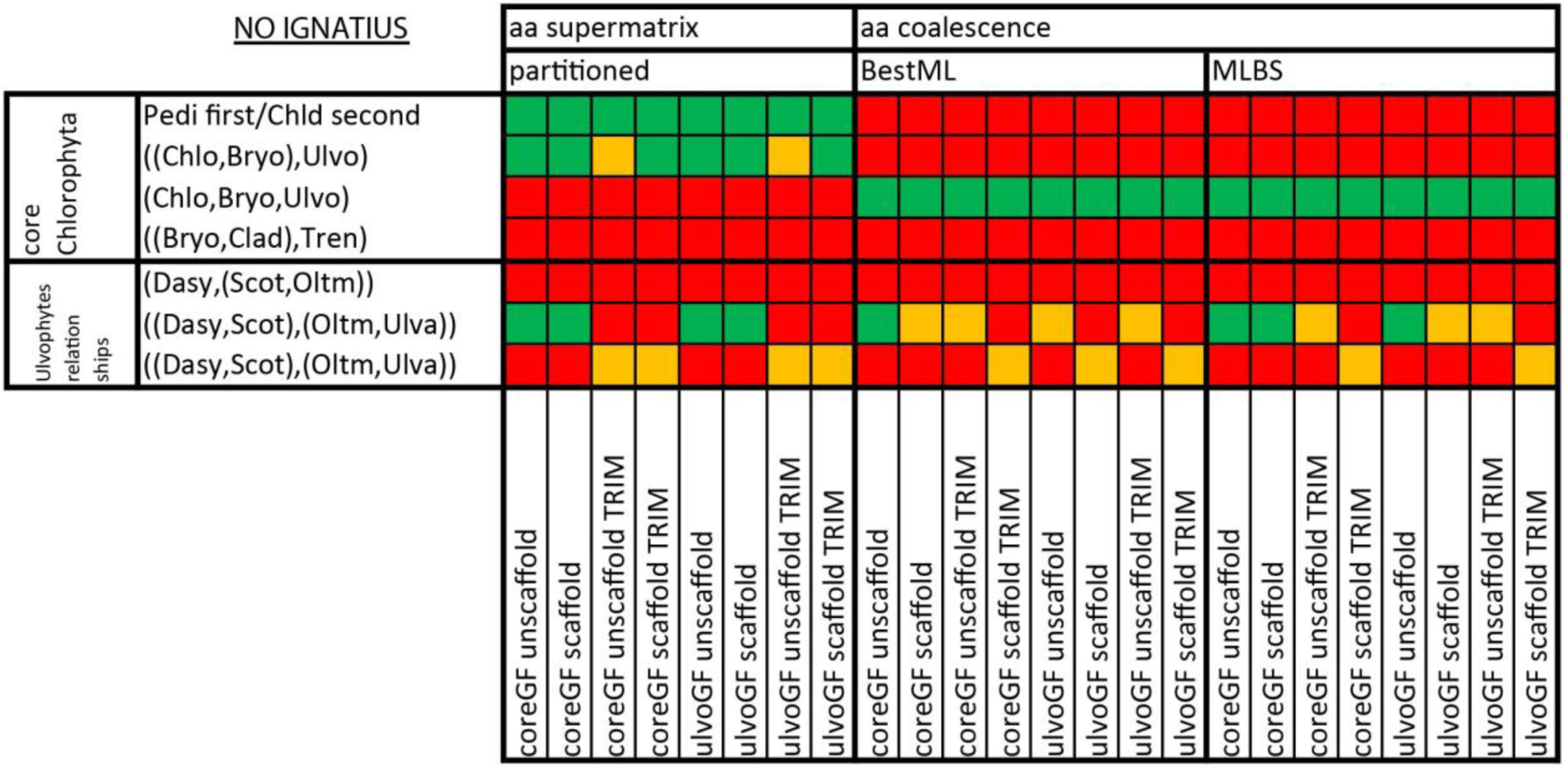
Summary of support for hypotheses of Chlorophyta relationships in analyses where *Ignatius* was pruned from the dataset. See legend of Figure S2 for details.

**Figure S6.**
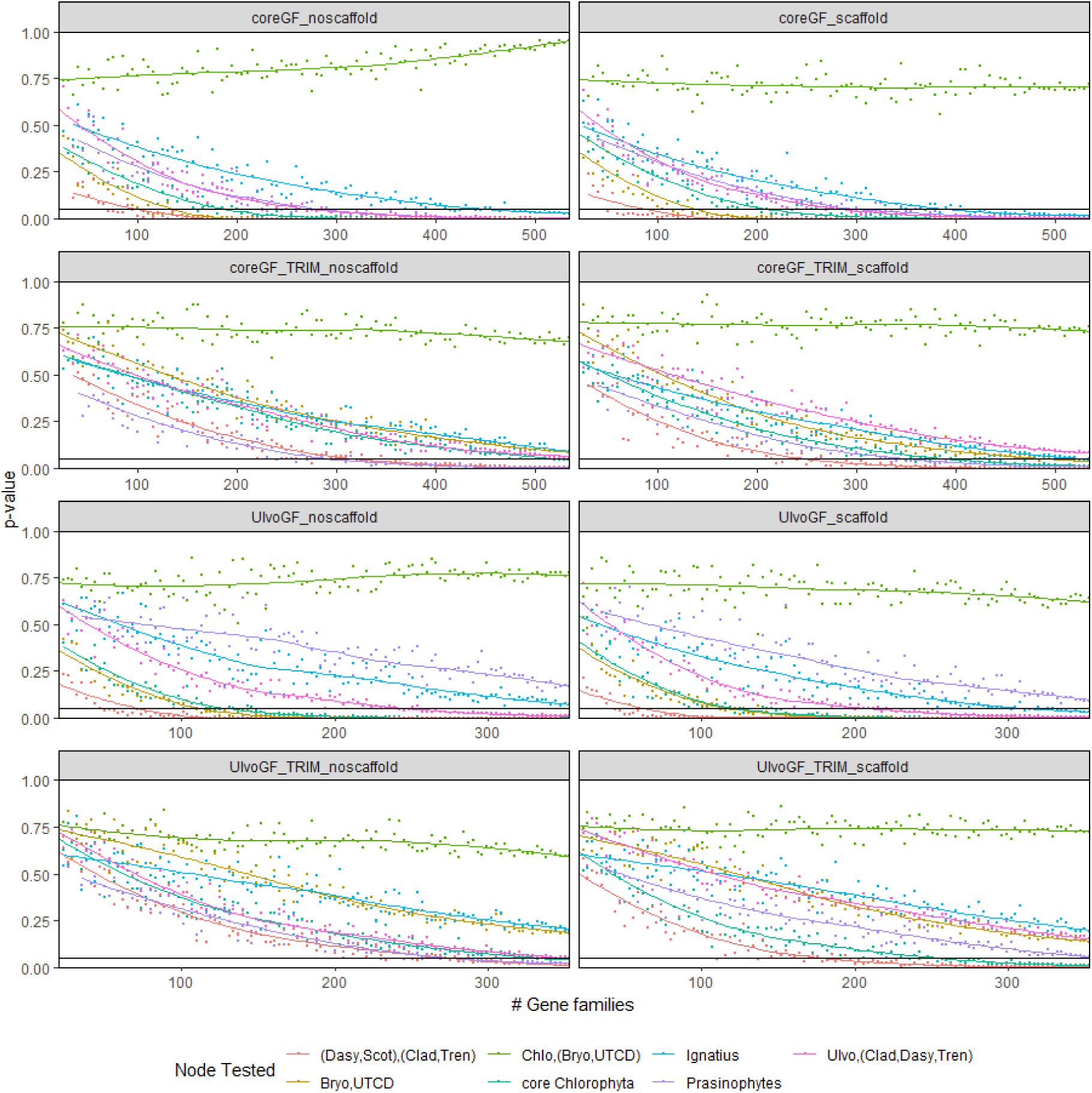
Tests for polytomy null-hypothesis. Polytomy test results for selected branches for each dataset. Gene trees were built with random growing subsets of genes (1%, 2%, … 100%, but not less than 20 genes), 10 replicates were run for each dataset and each subset. The median of the p-values of selected branches for each subset is plotted on the y-axis, against the number of genes in the subset (x-axis). The horizontal black line indicates p-value = 0.05.

**Figure S7.**
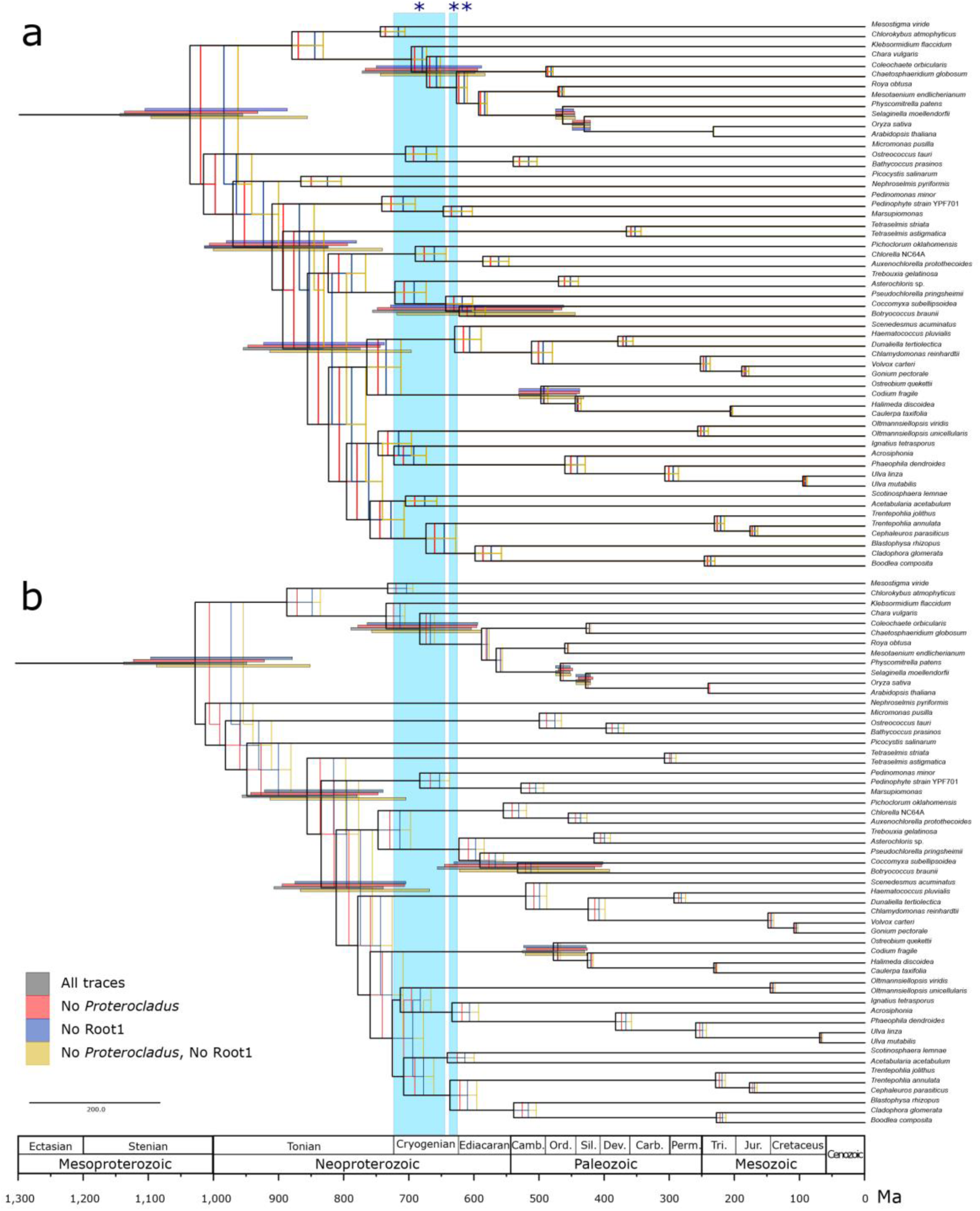
Summary of time calibrated phylogenetic trees based on different analyses, including all traces, analyses without R1 root, without the *Proterocladus* calibrating point, and without R1 root nor *Proterocladus* (Table S5, S6). Chronograms are based on the 10 most clock-like genes from the coreGF scaffolded trimmed dataset (see methods). Figure A shows the analyses constrained with the ML-based topology (T1), Figure B shows the analyses constrained with the coalescence-based topology (T4). Node values indicate average node ages. The bars represent 95% confidence interval (CI) of the calibration nodes relevant for this study. The two blue bars indicate the Sturtian (*) and the Marinoan (**) glaciations.

**Figure S8.**
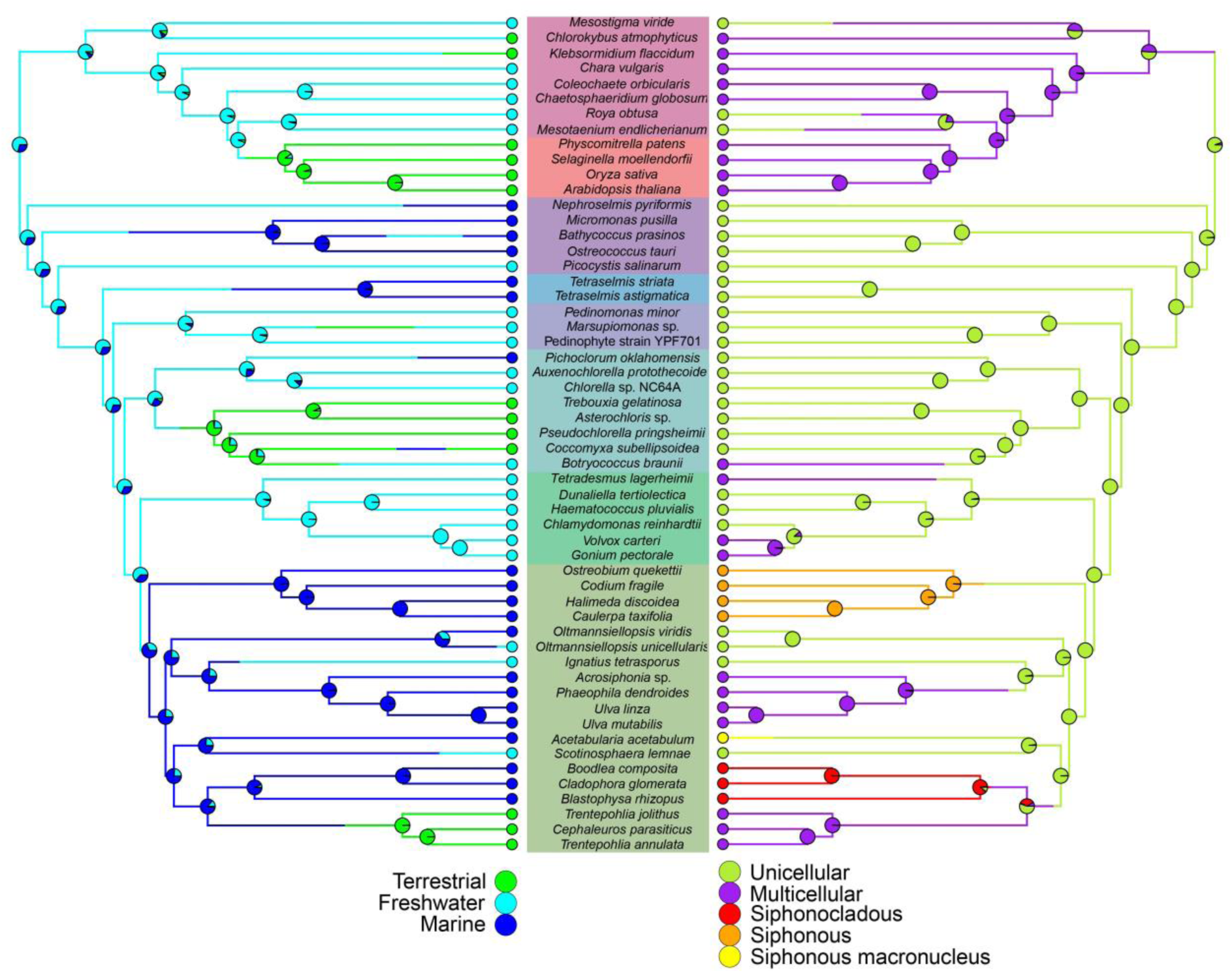
Ancestral state estimation of habitat (left) and cyto-morphological (right) traits, plotted on the ultrametric tree.

